# Microbial communities associated with marine sponges from diverse geographic locations harbour biosynthetic novelty

**DOI:** 10.1101/2024.01.09.574914

**Authors:** Vincent V Nowak, Peng Hou, Jeremy G Owen

## Abstract

Marine sponges are a prolific source of biologically active small molecules, many of which originate from sponge-associated microbes. Identifying the producing microbes is a key challenge in developing sustainable routes for production and isolation of sponge-associated metabolites, and requires application of several computational tools. To facilitate these analyses, we developed MetaSing, a reproducible singularity-based pipeline for assembly, identification of high quality metagenome-assembled genomes (MAGs), and analysis biosynthetic gene clusters (BGCs) from metagenomic short read data. We apply this pipeline to metagenome datasets from 16 marine sponges collected from New Zealand, Tonga and the Mediterranean Sea. Our analysis yielded 643 MAGs representing 510 species. Of the 2,670 BGCs identified across all samples, 70.8% were linked to a MAG, enabling taxonomic characterisation. Further comparison of BGCs to those identified from previously sequenced microbes revealed high biosynthetic novelty in variety of underexplored phyla including Poribacteria, Acidobacteriota and Dadabacteria. Alongside the observation that each sample contains unique biosynthetic potential, this holds great promise for natural product discovery and for furthering the understanding of different sponge holobionts.

## Introduction

The marine environment has long yielded interesting novel natural product chemistry with a variety of biological activities.^1,2^ In 2022, 20 marine natural products or derivatives from variety of chemical classes were in clinical use.^1,3^ Among marine invertebrates, sponges have been the most chemically diverse source of natural products, accounting for almost half of all marine natural products discovered.^2,3^ The capacity to produce secondary metabolites can be assessed by identification and annotation of biosynthetic gene clusters (BGCs) within microbial genome and metagenome data. BGCs are groups of co-localised genes that collectively act to produce small molecules that confer some selective advantage to their producer. While conclusively linking a natural product to its producing organism requires experimental validation,^4–7^ the bioinformatic analyses themselves are time-consuming and may require a significant time investment for researchers to acquire the necessary expertise.^8–10^

Metagenomics is the simultaneous analysis of a mixture of microbes, often via DNA-sequencing, and is widely used to study the unculturable majority of bacteria by targeting DNA directly isolated from the environment.^11–14^ Despite advances in sequencing technology and metagenomic assembly algorithms^8,15,16^ the reconstruction of resolved genomes on a single contig remains challenging and a process called binning is typically required to resolve individual genomes.^12^ A single binning algorithm may not produce the best results and two or more binning algorithms can be run in parallel followed by dereplication to select the best version of each bin from the different sets, with the final version referred to as a metagenome assembled genome (MAG).^9,17,18^ No single optimal workflow or combination of bioinformatic tools exist for the metagenomic assembly problem but recent pipelines and large-scale metagenomic studies show that the process of assembly, binning and bin dereplication generally produces the most informative and accurate results.^9,16,19–23^

Metagenomics has been immensely useful in microbial ecology and host-microbe studies but has also accelerated the identification of unprecedented secondary metabolic potential.^15,24,25^ Many marine sponge metagenomics studies focus on the overall metabolic capacity ^26–29^ relying on algorithms that identify clusters of orthologous groups of proteins (COGs) and/or Enzyme Commission numbers (ECs) that relate to biochemical pathways in the Kyoto Encyclopaedia of Genes and Genomes (KEGG) database.^30^ While these studies have been immensely useful for the study of the sponge holobiont ^31^ some are lacking analysis of secondary metabolism and many do not use specialized software needed to identify BGCs and the putative natural products produced by them.^32–35^ Here, we developed a reproducible singularity-based metagenomic pipeline that produces high quality MAGs, uses specialized software to identify BGCs and then links BGCs to the producing MAGs. We then applied this pipeline to 16 marine sponge metagenomes from various geographic regions including previously studied sponges from the Mediterranean sea ^36^ and New Zealand ^37^, as well as newly sequenced sponges from New Zealand and Tonga. This work reveals the taxonomic groups responsible for secondary metabolite production, demonstrates the novelty of some of the BGCs and the presence of vast biosynthetic potential in abundant members of the sponge microbiome.

## Results

### Development and testing of the MetaSing pipeline

Bioinformatic pipelines for metagenomic datasets have only recently become more popular and user-friendly. Thus far, they have been conda-based, like metaWRAP ^9^, Snakemake-based, like ATLAS ^38^ or MetaLAFFA ^39^ and Nextflow-based, like MUFFIN ^40^ but are largely focused on identifying general metabolic potential rather than specifically the biosynthetic potential for secondary metabolites and do not make use of the reproducibility offered by singularity containers. ^10,41^To address this, we developed the MetaSing pipeline, which follows the standard metagenomic workflow of trim-assemble-bin but utilizes three different binning algorithms and bin consolidation to achieve high quality MAGs.^9,17,18^ Assembly is achieved using metaSPAdes^42^, a high-performance metagenomic assembler ^15,16^ used in other marine sponge metagenomic studies.^22,28,37,43,44^ Binning tools used are metaBAT2 ^45^ and maxbin2 ^46^, two of the most widely used and tested binning tools ^8,16,18,21,28,37^, as well as autometa.^47^ Autometa is especially suited for microbiome analyses as it separates contigs based on Kingdom-level taxonomy and only uses bacterial contigs for binning, which significantly reduces host contamination.^21,37,47^ Following bin dereplication with dRep v.2.6.2 ^17^ to construct the final set of MAGs and their taxonomic identification using GTDB-Tk v.1.1.1 ^48,49^, BGCs are then linked to taxonomic groups, which to the best of our knowledge is currently not available as a pipeline. The pipeline is split over five singularity containers, which carry out separate steps of the analysis (Figure 1 and https://github.com/VincentNowak/meta_sing for more details).

**Figure 1.**
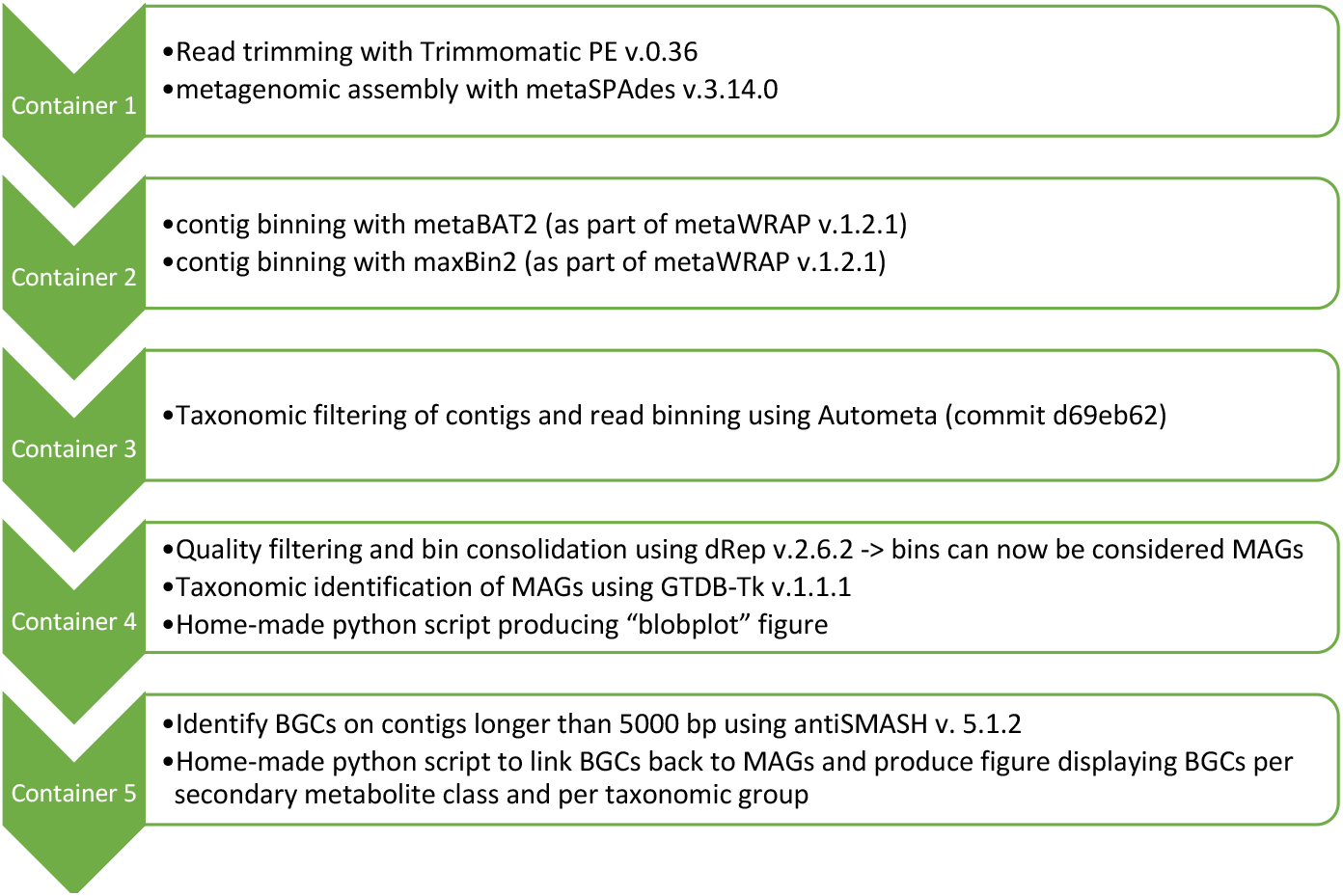
overview of the MetSing pipeline for rapidly identifying BGCs from metagenomic datasets and linking them to the producing MAG

Visual outputs provided by the MetaSing pipeline include differential coverage plots inspired by Albertsen and others (Figure 2A) ^50^ and a summary of BGCs by taxonomic group in the form of a collection of bar graphs (Figure 2B). Summary of results from MetaSing is given in Table S3 while data frames and figures created for each of the 16 marine sponge microbiomes are available on GitHub: (https://github.com/VincentNowak/PhD_thesis/tree/main/Chapter_2)

**Figure 2.**
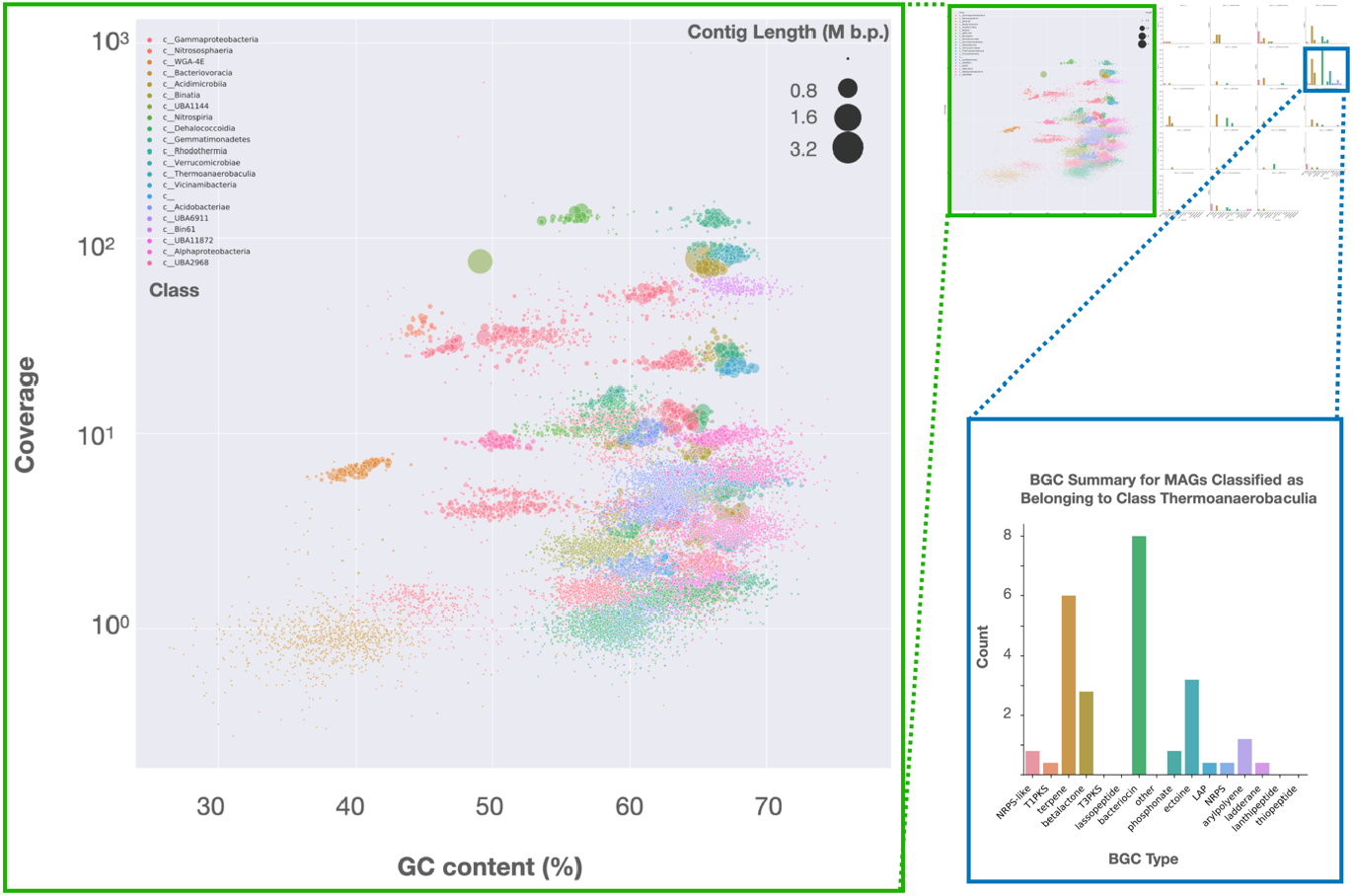
Screen shots of MAG summary plots produced by MetaSing: Data shown is for the MNP0977 sponge microbiome. Upper right thumbnail shows the entire layout for the respective graphs. A detailed view of the blobplot ^50^ and the BGC summary for one class from the Acidobacteriota is given.

### Genome-resolved metagenomics of 16 marine sponges

16 marine sponge metagenomes from a variety of geographical locations and covering several taxonomic groups within the Porifera (Table S1, S2) were analysed using the MetaSing pipeline. 643 MAGs were identified across all samples with the largest number (*n* = 80) originating from the deeply sequenced Mediterranean sponge *A. aerophoba* closely followed by 79 MAGs from the Tongan sponge CS200 (Table S3, Figure 3). Two NZ sponges, *M. hentscheli* and the MNP7375, a putative Halichondrida sponge, had much less complex microbial communities associated with them, as previously reported for *M. hentscheli*.^37,43^ The average completeness of all 643 MAGs is 89.5% and average contamination is 3.3%, which is comparable to other sponge-specific datasets.^22,28^ 286 of the 643 MAGs can be considered high quality being at least 90% complete with less than 10% contamination.^49,51–53^

**Figure 3.**
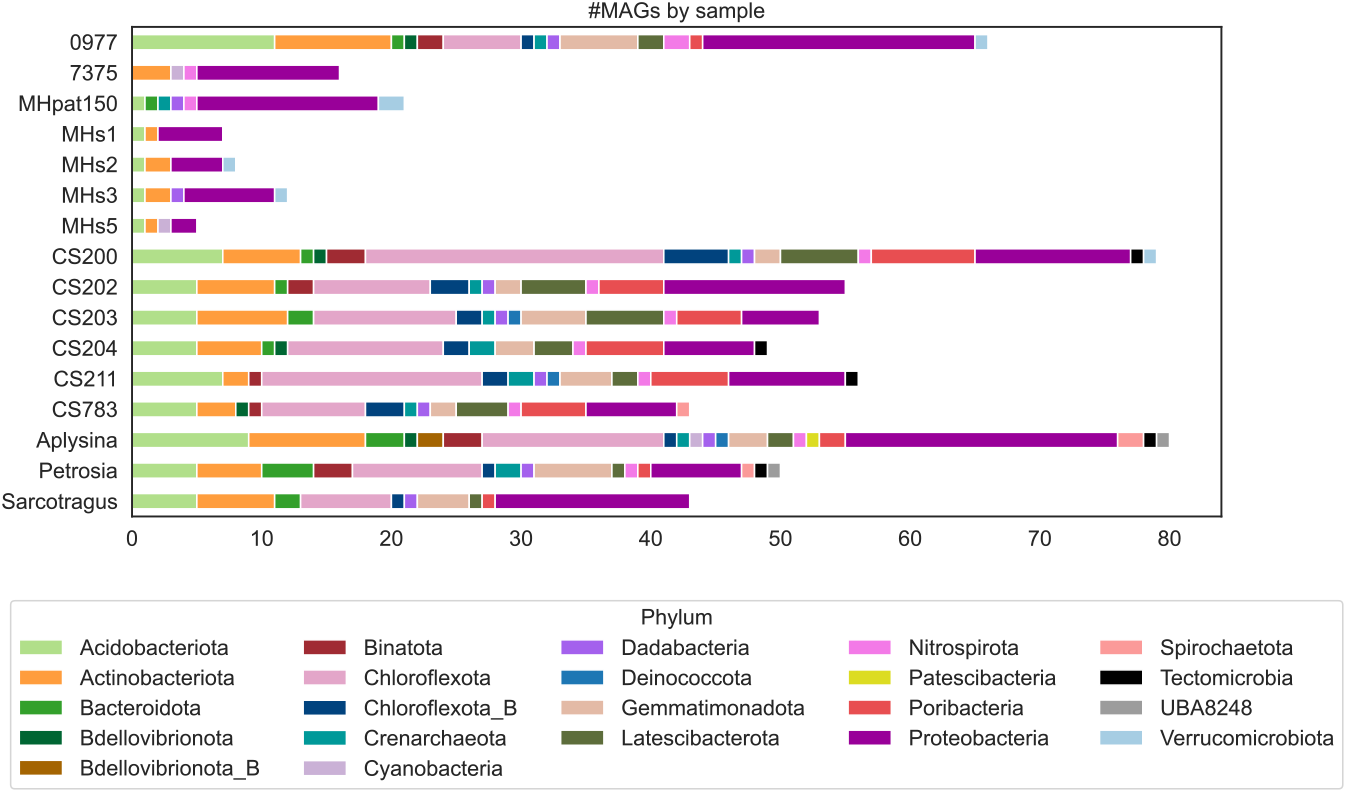
Deduced microbiome composition for each sample in this study: Number of MAGs by sample and phylum level taxonomy are shown. Taxonomy was assigned to MAGs using GDBTK.

In total, 22 different phyla were identified by GTDB-Tk ^49^ across all 16 marine sponge metagenomes (Figure 3). The three most commonly occurring phyla were Proteobacteria, Chloroflexota, and Acidobacteriota (Figure S1). These microbial phyla are consistent with the phyla identified in other sponge microbiome studies.^22,28,54^ Notable was a large number of Poribacteria and Latescibacterota MAGs in the Tongan sponges (Figure 3). Latescibacterota include the PAUC34f, which was previously designated a Candidate phylum and originally discovered in sponges, but have recently expanded to include other lineages mostly from the marine environment.^55^ Taxonomic classification using GTDB-Tk as part of the MetaSing pipeline resulted in all MAGs being classified at the phylum level, all but 8 MAGs classified at the class level and all but 52 MAGs classified at the order level. The 8 MAGs not classified at the class level all belong to the Latescibacterota, indicating their phylogenetic novelty compared to reported members of this phylum (Figure S2A). Of the MAGs not classified at the order level, most (*n* = 14) belonged to the Acidobacteriota (Figure S2B), highlighting the phylogenetic diversity added by this work.

### Comparative metagenomics and abundant community members

Host taxonomy and the surrounding environment have been identified as one the main factors to influence microbiome composition but composition can also vary between time points or DNA extraction protocols ^27,36,56^ and intraspecific variation may be higher in some sponge species.^57–59^ Dereplication of the 643 MAGs using dRep v.2.6.2 with 0.95 as the ANI cutoff ^19,22,28,49,60^ resulted in 510 unique “species-level” MAGs (hence forth referred to as species). As expected numerous MAGs (*n* =26) were present in multiple *M. hentscheli* samples. In contrast, MNP0977 did not have any MAGs that were found in other metagenomes, which indicates this sponge contains unique microbial species not found in the other sponges. The overall taxonomic trends and dominant phyla remained largely the same at the species level (Table S4). Out of the abundant phyla, the largest changes were seen in the Acidobacteriota, Poribacteria and Latescibacterota, which indicates many shared members across different samples and/or large strain diversity.

In order to examine the influence of geography and sponge taxonomy on microbiome composition, reads from each of the 16 datasets were mapped against the 510 species to quantify their abundance and allow comparison of the sponge metagenomes. The 16 datasets were clustered using both binary presence/absence (Figure 4) and continuous relative abundance values obtained from read mapping (Figure S3). Simple binary presence/absence clustering was included to reduce the potential effects of stochastic variation in the abundance of microbiome members, which may occur between different specimens or time points.^27,28,31,36,57^ As expected, the five *M. hentscheli* samples clustered together with both binary and continuous clustering. An unexpected result was the grouping of MNP7375 (putatively a *Halichondria* species) with *M. hentscheli* samples indicating a similar microbiome composition albeit with differing relative abundances of species. The sponge microbiomes examined in this work consistently cluster by geographical region using binary clustering with the MNP0977 (putatively a Irciniidae species) distantly related to other NZ sponges (Figure 4). When using continuous clustering the clustering by geographic region was less apparent (Figure S3), which highlights the utility of using binary clustering to mitigate stochastic variation when comparing microbiomes. Collectively, these analyses support the idea that both geographic region and host taxonomy have an influence on the sponge microbiome.

**Figure 4.**
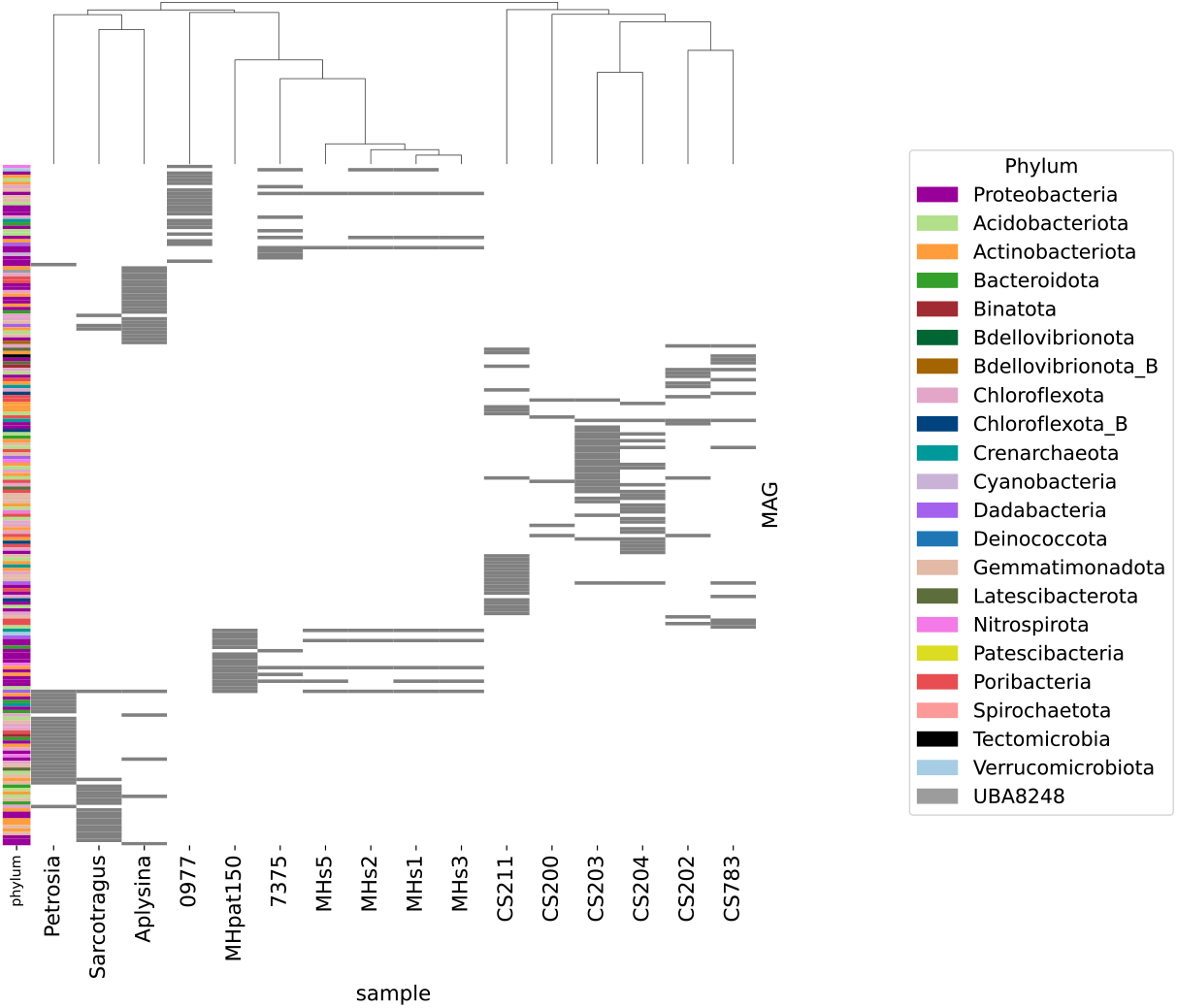
Comparison of microbiome composition across sponge samples. Figure shows binary presence/absence of all MAGs with a relative abundance > 1% in any of the samples. Samples are clustered by cosine distance using complete linkage

Closer investigation of the 10 most abundant MAGs in each sample (Figure 5) revealed that the microbiomes of some sponges were dominated by a single member (CS200, CS783), whereas others had a relatively even abundance of several members (Aplysina, Petrosia, MNP0977). High abundance of a single member does not appear to be correlated to the use of PCR during sequencing library construction (Table S1). While it is unclear whether these abundance patterns are due to temporal variation or more permanent, it is evident that a single or few microbe(s) may dominate the biomass within a sponge at a given timepoint. This poses the question of what functional roles these microbes may be fulfilling at the sampling timepoint and in particular, whether these may be related to the production of defence chemicals as observed in *Theonella swinhoei* ^6^, *Lamellodysidea herbacea* and *Dysidea granulosa* ^4^, and *Haliclona* sp. ^44^, or are related to primary metabolic interactions. For example, a single species of Crenarchaeota is abundant in 10 of the 16 samples, all of which are from NZ or Tonga. Host-associated archaea, such as may this species of Crenarchaeota, may fulfil a suite of functions but most commonly carry out methanogenesis ultimately supporting bacterial and protist metabolism.^59^ High abundance of Crenarchaeota has been reported in some sponges in the Global Sponge Microbiome based on 16S rRNA amplicon sequencing ^54^ and they are putative keystone members of the *Ircinia ramosa* microbiome governing the first step in nitrification.^28^ A Poribacteria species identified as *Candidatus* genus WGA-3G has a very high abundance (∼75%) in CS200 but a low abundance (<1.6%) in CS202, and CS211 (all three putative Dictyoceratida species) as well as CS203 and CS204 (putative Verongiida species). This taxon was first reported in a single cell whole genome amplification study as encoding several important enzymes for primary metabolism including sulfatases, sugar transporters, uronic acid degrading enzymes and enzymes involved in glycolysis.^61^

**Figure 5.**
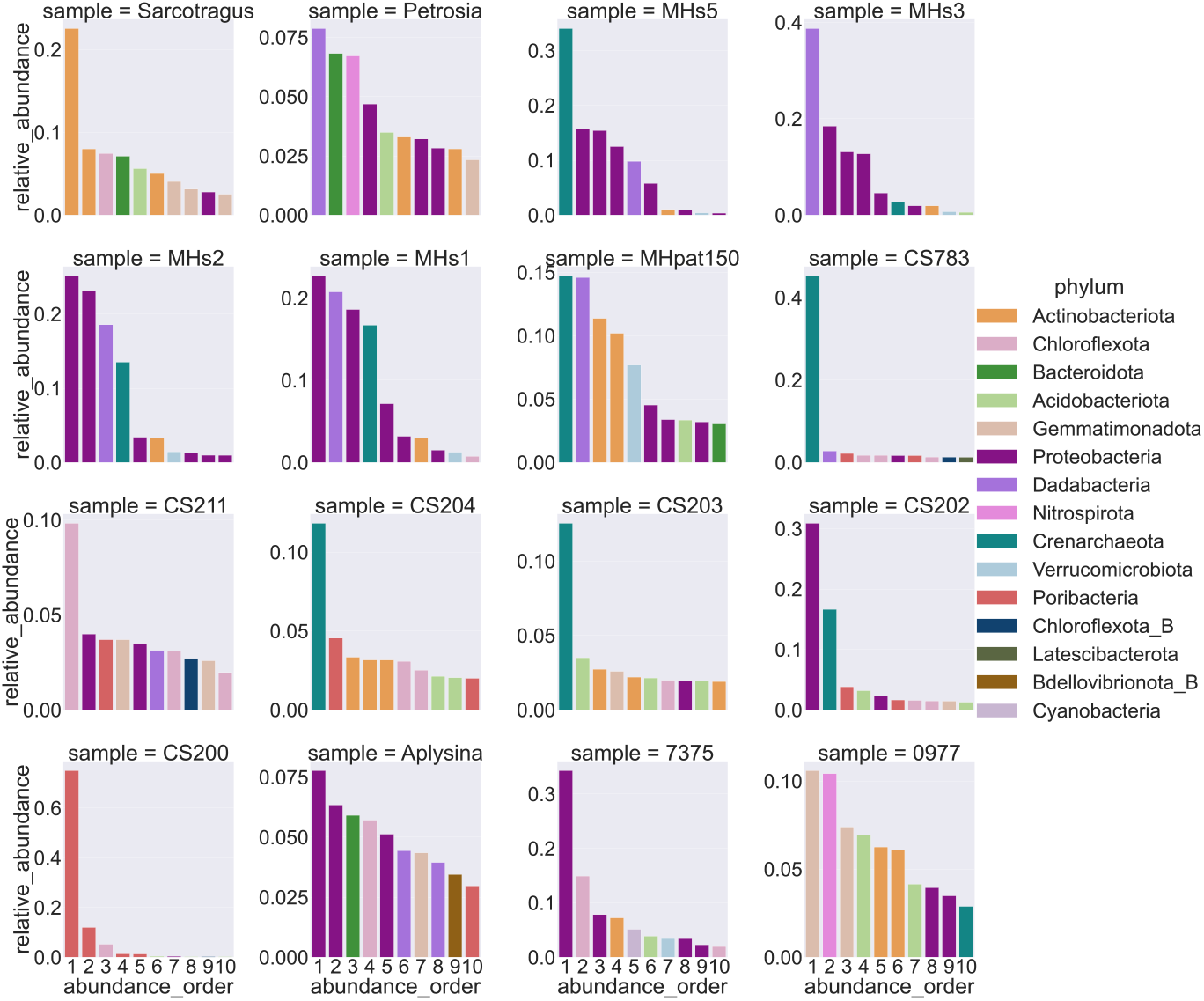
Relative abundance of the 10 most abundant MAGs in order for each sample in this study.

### Biosynthetic gene cluster diversity

A total of 2,670 BGCs were identified from the 16 sponge metagenome assemblies (Table S3), 70.8% (*n* = 1,890) of which were linked to a MAG and 28.7% (*n* = 767) of which can be considered complete (not on contig edge). BGC here refers to the whole region identified by antiSMASH5 since detailed comparison and experimental work would be required to determine if a region contains one or several distinct biosynthetic pathways.^62–64^ The resulting 52 product types predicted by antiSMASH were summarised into 11 broader BGC classes for simplicity (Table S5). Across the 16 metagenomes terpenes were the most commonly identified BGC followed by PKSs (especially Type 1 PKSs), bacteriocins and NRPSs (Figure S4). Few BGCs >50 kb were assembled overall (Figure S5). BGC fragmentation did not appear to be due to insufficient sequencing depth as these factors did not correlate well (Figure S6), and it is likely that this occurred due to repeat regions commonly found in BGCs, such as those in NRPSs and PKS BGCs known to result in fragmented assemblies.^37,43,53,62^

The largest number of terpene and bacteriocin BGCs was attributed to Proteobacteria, which also account for most of the MAGs identified across all samples (Figure S1, S7). While relatively equal number of PKS BGCs was identified from Proteobacteria and Acidobacteriota, the latter accounted for more NRPS BGCs (Figure S7, S8). Acidobacteriota also account for the highest number of RiPP, RiPP hybrid and bacteriocin-RiPP hybrid BGCs despite being not as common as Proteobacteria. The prominence of diverse secondary metabolism across the Acidobacteriota identified in this work is evident at the MAG-level with a large number and variety of BGCs per MAG across the phylum (Figure 6A, Table S6). Verrucomicrobiota on the other hand have the highest number of BGCs per MAG but all BGCs are fragmented, reflecting the fact that these are predominantly large NRPS and PKS BGCs. Variability in the number and type of BGCs in MAGs within the same dRep species cluster (Figure 6A, Figure S9) may occur due to varying assembly and binning qualities across samples but may also be due to strain-level differences in BGCs as reported for the *Salinispora* ^65,66^ and *Streptomyces* ^67^ among others.^68^ Using the percentage of the genome covered by BGCs (BGC %) as another metric of biosynthetic richness, the top three phyla were Verrucomicrobiota, Nitrospirota and Acidobacteriota (Table S7).

**Figure 6.**
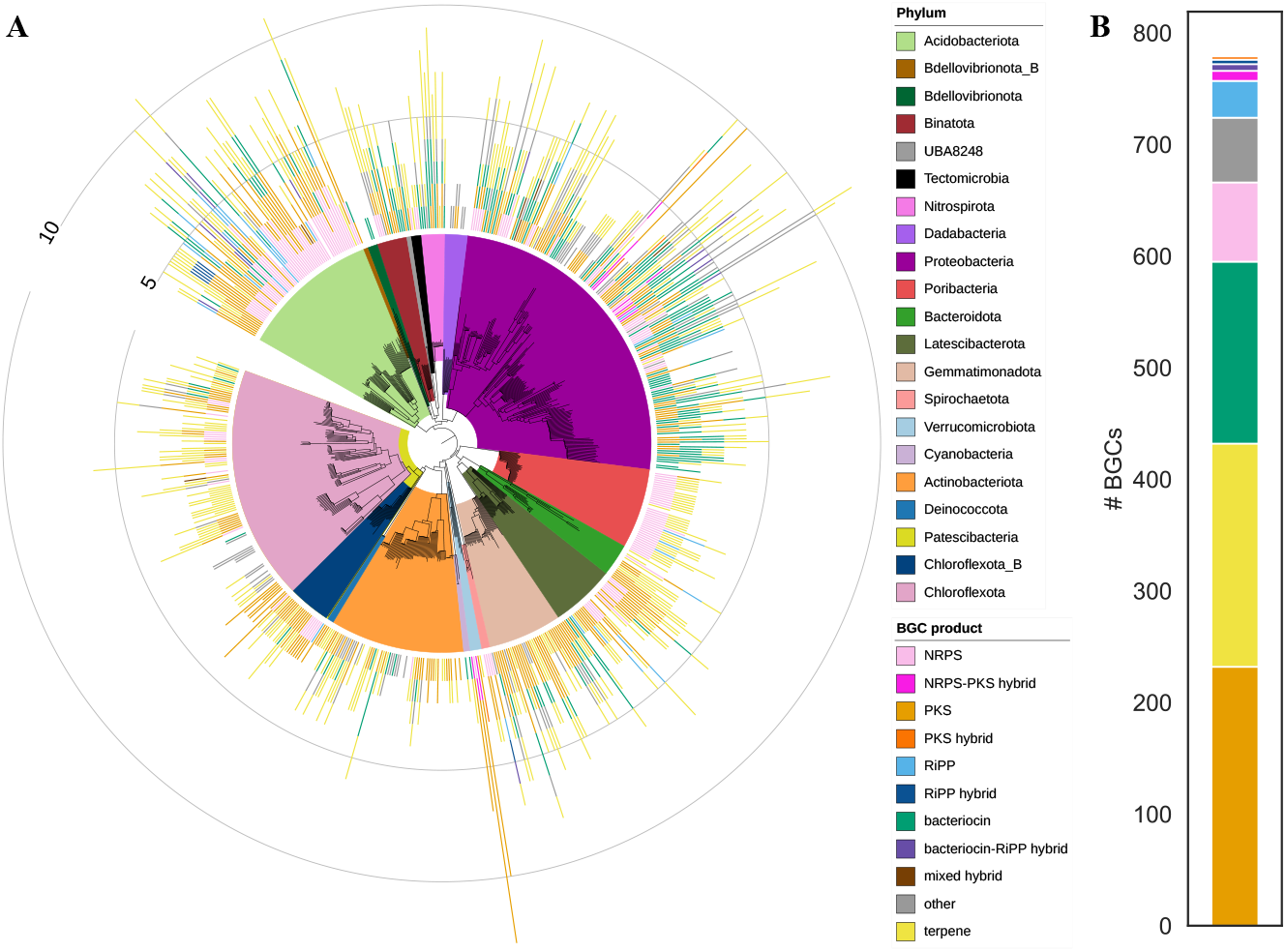
Biosynthetic characteristic of all MAGs identified in this study: **A** Phylogenetic tree of all 643 MAGs coloured by phylum, showing the number of BGCs per BGC product class (summarised as outlined in Table S5) per MAG (bars radiating from centre). Figure was created using iTol ^308^ and data for interactive exploration is available on GitHub (see 2.2.5). Figure S9 shows the phylogenetic tree in more detail as well as sample, region and dRep clusters **B** Number of BGCs not associated with a MAG (*n* = 780)

### Clustering of BGCs to infer relatedness and novelty

In order to identify biosynthetic novelty all BGCs identified in this work were queried against ∼1.2 million BGCs using the query function of BiG-SLICE.^69^ Analysis using BiG-SLICE aims to identify related BGCs based on vectorization of BGC features/domains and their Bit-scores, which are queried against profile Hidden Markov Models to derive a centroid vector (akin to a consensus BGC), against which the individual BGCs are then queried again to form so-called gene cluster families (GCFs). GCFs were created using complete BGCs (*n* = 802,287) initially and incomplete BGCs were then queried back onto these GCFs for the final network. Following this workflow, complete and incomplete BGCs identified in this work were queried against the BiG-SLICE model using the more conservative T=900 threshold, which is expected to cluster even distantly related BGCs.^69^

The membership value (*d*) produced by BiG-SLICE during GCF creation gives an indication of how confidently a BGC is assigned to a GCF where *d* ≤ T are “core”, T < *d* ≤ 2T are “putative” and *d* > 2T are “orphan” members. The 2,670 BGCs in our analysis were attributed to 255 GCFs, where 59 BGCs (2.20%) are orphan members (Figure S10A, S10B). Several BGCs were attributed to very small GCFs and closer inspection using clinker ^70^ showed that BGC similarity was very low with very few genes shared. This is highlighted by GCF_08405, which contained Type I PKS BGCs specific to putatively different species of Latescibacterota in Tongan sponges (Figure S10C, S10D). The MiBIG database ^71,72^ is a database containing experimentally characterised BGCs and was included in the BiG-SLICE model. Of the 225 GCFs that BGCs from this work were attributed to, 153 did not contain MiBIG BGCs, accounting for 936 (35.06%) of the BGCs queried. This, in concert with the observed low similarity within GCFs created by BiG-SLICE, suggests a potentially high biosynthetic and chemical novelty.

Recent studies have shown that improvements to the BIRCH clustering algorithm used in BiG-SLICE ^73^ or the use of cosine distance ^23,74^ to cluster BGCs can remediate some of the observed inaccuracies BiG-SLICE, especially for shorter BGCs with fewer features. To this end, the minimum cosine distance of all 2,670 BGCs relative to the BiG-FAM database ^69^ inherent to BiG-SLICE was calculated. BGCs with a minimum cosine distance (min(d_BiG-FAM_)) larger than 0.2 were considered novel as previously determined.^23^ Using cosine distance appeared to better characterise novelty of shorter BGCs, such as RiPPs and bacteriocins, while maintaining the distribution of most other major natural product classes (Figure 7A, Figure S10A). 56.22% (*n* = 1,501) of BGCs were identified as novel and included BGCs from all major classes of natural products. This includes the above-mentioned GCF_08405 comprised of Type 1 PKS BGCs associated with Latescibacteriota, and numerous of the previously identified sponge-derived RiPP proteusins ^75^, which were identified in a variety of bacterial phyla (Figure S11). The highest novelty rate (# novel BGCs/# BGCs) was observed in hybrid BGC classes like RiPP hybrid, mixed hybrid and RiPP-bacteriocin hybrid as well as the Other and NRPS classes (Table S8). Notably, the NRPS and Other BGC classes have an even distribution of numerous BGCs across the whole range of min(d_BiG-FAM_) values, indicating a trove of biosynthetic novelty to be discovered (Figure 7A).

**Figure 7.**
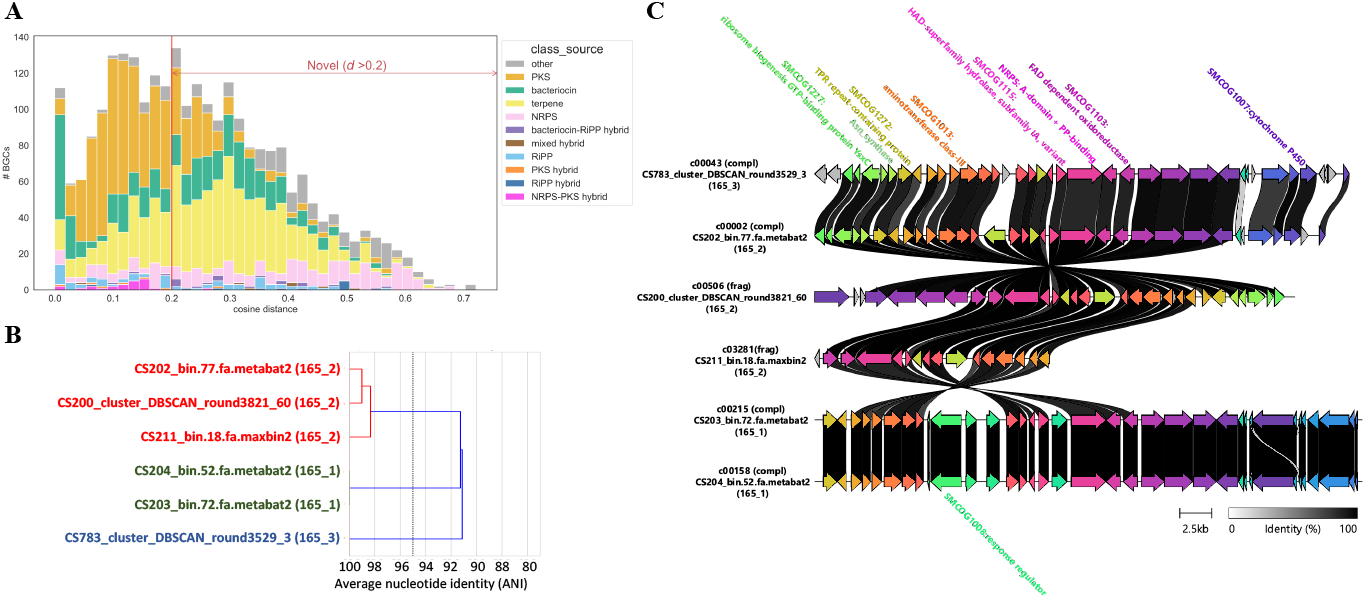
Marine sponges harbour unusual biosynthetic gene clusters: **A** Histogram of cosine distance (min(d_BiG-FAM_)) for all 2,670 BGCs identified in this study with colour indicating the natural product class (see Table S5 on how these were summarised) **B** dRep secondary cluster containing closely related species (165_1, 165_2 and 165_3) from the Acidobacteriota **C** GCF_245 containing conserved NRPS BGCs identified from all six Tongan sponges, with (frag) indicating an incomplete BGC and (compl) indicating a complete BGC. Only annotations given by antiSMASH are given for visibility (see Table S10 for supplementary BLAST results).

The highest number of novel NRPS BGCs associated with a phylum were from the Acidobacteriota (*n* = 80) and the Poribacteria (*n* = 35). Several of these BGCs were the only BGC in the respective GCF but some GCFs with high novelty were present in several samples. Indeed, a highly conserved GCF present in closely related species of Acidobacteriota could be found in all six Tongan sponges (Figure 7B, 7C), indicating that the secondary metabolite produced by this non-canonical NRPS may have an evolutionary advantageous function for these Tongan sponge holobionts. The MAG which contains one of these BGCs, is a cryptic MAG identified only at the class level as belonging to the Acidobacteriae and is also among the 10 most abundant MAGs in the CS203 sample (Figure 5, 7C). In fact, the MAG with the highest number of novel BGCs (*n* = 8), a putatively novel genus of Acidobacterium in the UBA8438 family, is also among 10 most abundant MAGs (Figure 5). Prediction of the compound(s) produced by these NRPS BGCs is difficult due to the non-canonical nature of this Type III NRPS ^76^ but appears to be a highly modified peptide containing methylated, acetylated, hydrolysed and otherwise modified amino acid residues. Poribacteria have been well-studied in the context of the sponge holobiont and have been associated with a variety of functions but largely focused on nutrient exchange and primary metabolism. ^61,77^ Similar to the Acidobacteriota, GCFs with high novelty associated with Poribacteria were dominated by NRPSs and included singleton BGCs as well as BGCs shared across samples, often specific to the Tongan sponges. GCF_18, for example, contains Type III NRPS BGCs that appear to incorporate several non-proteinogenic amino acids including arginosuccinate and 8(S)-amino-7-oxononanoate, and are found in a single species of the Poribacteria present in four of the six Tongan sponges (Figure S12). Other phyla with a high rate of novel BGCs but a smaller total number of BGCs include the Tectomicrobia, which are known producers of secondary metabolites ^7^, Deinococcota, Dadabacteria and Spirochaetota (Table S9, Figure S14). While the above-mentioned high novelty GCFs are shared within a region, the largest number of shared GCFs is between all Mediterranean and all Tongan sponges (Figure S13). Analysis of shared GCFs also highlights the large number of sample-specific GCFs, which indicates that each marine sponge sample harbours unique biosynthetic potential.

## Discussion

Evidence is growing that various phyla associated with marine sponges produce bioactive compounds. Since the discovery of the *Entotheonella* genus from the *Theonella swinhoei* sponge ^5,7^, Cyanobacteria ^4^, Proteobacteria ^44^, Verrucomicrobia ^78^ and Chloroflexota ^79^ have been conclusively linked to the production of natural products in marine sponges. Here, we present MetaSing, a reproducible bioinformatic pipeline that allows facile identification of sponge-associated taxa encoding biosynthetic potential. While, this study demonstrates the utility of MetaSing in studying sponge microbiomes, it may in principle be applied to other short read metagenomic datasets. Using MetaSing, we demonstrate that various different phyla associated with marine sponges are rich in BGCs, and that these phyla may differ between host species. Acidobacteriota for example, which are known to harbour a considerable number of BGCs in soil ^80^, stood out in this and other work ^81^ as containing biosynthetic richness and novelty. Also noteworthy, is the biosynthetic novelty identified in the Poribacteria, which have long been known to be closely associated with marine sponges ^77^, and other phyla less commonly studied in the context of the sponge holobiont, like the Deinococcota, Dadabacteria and Spirochaetota. Alongside the evidence presented here that each sponge metagenome contains unique biosynthetic potential, this demonstrates that sequencing of new sponge metagenomes is likely to continue leading to the discovery of further biosynthetic potential and natural products.

Clustering of BGCs into GCFs and quantifying their novelty is a relatively new approach. The BiG-SLICE BGC network used as a reference is based on ∼209,000 RefSeq genomes but includes draft genomes and > 20,000 MAGs.^69^ Overall some phyla might still be underrepresented in this dataset, e.g. Dadabacteriota, and a certain database bias introduced into the novelty analyses. However, even for underrepresented taxa, the underlying truth that these BGCs may be novel still holds since the taxa themselves are novel or understudied. As touched on above, the use of Euclidean distance to compare BGCs and BIRCH clustering to form GCFs and centroid BGCs in BiG-SLICE allows rapid analysis of a large number of BGCs but compromises on accuracy compared to its predecessor BiG-SCAPE. ^69^ Improvements to the BIRCH clustering algorithm ^73^ or the use of cosine distance ^23,74^ appear to remediate some of the observed inaccuracies in BGC clustering, as we have found to be true for cosine distance in this study. Rather than focusing on all BGCs in the BiG-SLICE reference, some studies ^81,82^ focus on the characterised BGCs deposited in the MiBIG database to identify chemically uncharacterised space by those BGCs that do not cluster with MiBIG BGCs. In any case, the study of BGCs identified from metagenomes and comparisons thereof are subject to biases, like variations in DNA extraction, the use of PCR in the sequencing library construction protocol as well as temporal variation of the microbiome and the transient presence of some microbes ^27,36,56^ Metagenomic assemblies in general may be influenced by uneven coverage, strain heterogeneity and genomic repeat regions as well as the need for high coverage to resolve genomes.^17,50,83,84^ During binning, contigs, and thus BGCs, may not be attributed to the correct MAG or as is more commonly the case not be attributed to any MAG.^78^ While a BGC typically contains most or all of the biosynthetic enzymes, regulatory genes, resistance elements and transporters required for compound production,^85^ unusual genes or gene arrangements including the use of genes that are not co-localised may occur ^86,87^, which may further complicate correct attribution. The pipeline presented here allows facile processing of numerous datasets and facilitates selection of samples containing taxonomic groups and/or BGCs of interest based on relatively inexpensive shotgun sequencing. Samples of interest can then be investigated further using long-read sequencing or other techniques, such as cosmid library construction ^88^, single-cell sequencing or strain isolation.^89^

While this study identified unique biosynthetic potential in each sample, which holds promise for natural product discovery, common biosynthetic potential was also identified across almost all samples, which suggests conserved and evolutionarily advantageous functions. We specifically identify near-identical BGCs that occur in numerous sponge metagenomes. These Type III NRPSs are not only highly conserved, but also distinct from previously identified BGCs, and may produce yet-uncharacterised natural products via novel biosynthetic mechanisms. In some cases, these BGCs were attributed to highly abundant community members. While it is unclear if the high abundance of said bacteria in this study is due to compounds derived from these BGCs or due to other trophic interactions like nutrient exchange, other natural product producers like *Entotheonella* ^7,90^ or Candidatus Endohaliclona renieramycinifaciens ^44^ can be highly abundant. In conclusion this study provides a reproducible metagenomics pipeline, which could also be applied to metagenomic datasets not originating from sponges, highlights taxonomic groups encoding large numbers of BGCs and identifies BGCs, GCFs and taxa which may encode secondary metabolites essential to the respective sponge holobiont.

## Methods

### Sample collection and DNA extraction

All samples in this study were collected by SCUBA diving with details summarized in Table S1. New Zealand and Tongan sponges were frozen without preservatives after collection. Metagenomic DNA was extracted as described by Storey and others.^37^ In brief, sponge tissue was ground under liquid nitrogen and enriched for prokaryotic cells by differential centrifugation followed by extraction with a sponge lysis buffer (8 M urea, 2% sarkosyl, 1 M NaCl, 50 mM EDTA, 50 mM Tris-HCl, pH 7.5). DNA was then purified using phenol-chloroform-isoamyl alcohol followed by removal of polysaccharides using CTAB and DNA concentrated using an ethanol precipitation.^91^

### Acquisition of sequence data

For the New Zealand and Tongan sponges, purified metagenomic DNA was sent to Annorad for library preparation and sequencing. Library preparation was achieved using a TruSeq workflow and PE150bp reads acquired using an Illumina HiSeq4000 instrument resulting in producing 10.58 – 43.44 Gb per specimen. For the Mediterranean sponges, reads were retrieved from Sequencing Read Archive (SRA) using prefetch followed by fastq-dump -F —split-files —gzip from the SRA toolkit. Datasets for *Aplysina aerophoba* are SRR6214543, SRR8092555, SRR8093131, SRR8092709, SRR8092710, SRR8092756, while *Petrosia ficiformis* is SRR3473501 and *Sarcotragus foetidus* is SRR3473948.^36^ The reads from the five *Aplysina aerophoba* datasets were pooled prior to assembly since they are arise from different tissue layers of the same specimen.^36^

### The MetaSing pipeline

All 16 metagenomic sequencing samples were analysed with the MetaSing pipeline https://github.com/VincentNowak/meta_sing.Container_1 runs a custom script (adap_ID.sh, Matt Storey) to identify adapter sequences in metagenomic reads, which are then appended to the adapter file provided as part of Trimmomatic 0.36.^92^ Reads are then trimmed and quality-filtered using Trimmomatic v.0.36 followed by assembly with metaSPAdes v.3.14.0.^42^ Container_2 runs metaBAT2 ^45^ and maxbin2 ^46^ from the binning module of metaWRAP v.1.2.1 ^9^ to construct two sets of bins from the metaSPAdes assembly. Container_3 runs the binning algorithm autometa.^47^ Container 4 runs dRep v.2.6.2 ^17^ to quality-filter (≥ 75% complete, ≤ 25% contamination, ≥ 50 kb size) bins, as determined by checkM: a program that uses lineage-specific marker-genes to determine these metrics ^17,51^, followed by ANI-based comparison ^52,60^ to consolidate the three bin sets into a single set of MAGs. Final MAGs are then taxonomically identified using GTDB-Tk v.1.1.1 ^48,49^ and visualised using a python-script. Container_5 runs antiSMASH5.1.2 ^63^ to identify secondary metabolite BGCs from all contigs (≥ 5,000 bp) and maps BGCs to MAGs.

### Summary figures for all samples

Based on the ‘region concept’ introduced in antiSMASH5, a region may contain more than one so-called ‘candidate cluster’, which are one of four types (single, chemical hybrid, interleaved or neighbouring).^63^ The BGC product type used in summary figures here is for each region and was formed by concatenating the predicted product types of all candidate BGCs in the respective region. (https://github.com/VincentNowak/PhD_thesis/blob/main/Chapter_2/Final_BGC_summary_110223.ipynb). Figures were created using seaborn and matplotlib as visible in the Jupyter notebook. The iTol BGCs per MAG summary was created by first running GTDB-Tk v.1.1.1 (as above) on all 643 MAGs to create the phylogenetic tree, which was then edited in Dendroscope v.3.5.10 ^93^ and used as the input for the iTOL v.6.6 online platform ^94^ alongside data derived from the above-mentioned Jupyter Notebook. Files for interactive exploration of this data are available on GitHub: (https://github.com/VincentNowak/PhD_thesis/tree/main/Chapter_2/iTol_data).

### Read mapping analyses to quantify abundance

The 643 MAGs collectively identified from the 16 datasets were dereplicated using dRep ^17^ at a 95% ANI cutoff to delineate unique “species” and reduce the effect of reads mapping ambiguously to strains or species shared across samples. To accurately quantify the abundance of individual MAGs, raw metagenomic reads were simultaneously mapped to all 510 dereplicated MAGs for each of the 16 datasets using bbsplit.sh from BBTools v.38.31 (BBTools – Brian Bushnell - sourceforge.net/projects/bbmap/). Only contigs covered by at least 75% were used in the analysis to reduce potential mis-mapping. Relative abundance of MAGs within each sample was then calculated based on the average fold coverage of all contigs associated with a MAG. Due to the absence of sequencing data from surrounding seawater a conservative abundance cutoff of 1% relative abundance ^57^ was used to define the core community of the sponge holobiont. In order to provide a qualitative overview of similarity between the metagenomes, vectors of relative abundance and binary presence/absence for MAGs in each metagenome were then clustered using the seaborn v.0.11.2 clustermap function (hierarchical clustering, cosine distance, average linkage). Cosine distance calculates the distance between vectors, making this a simple but effective metric for comparison that has been used in other genetic comparisons.^23,95^ The workflow is available as a Jupyter notebook on GitHub: (https://github.com/VincentNowak/PhD_thesis/blob/main/Chapter_2/Sample_vs_abundance_table_131022.ipynb).

### BiG-SLICE and cosine analyses

All 2,670 BGCs collectively identified from the 16 datasets were analysed using BiG-SLICE v.1.1.0 in --query mode to query BGCs against the database of ∼1.2 million BGCs created in the paper.^69^ The BiG-SLICE output was then manually inspected using the online server and summarised in a Jupyter notebook using the python sqlite3 library as detailed on GitHub (https://github.com/VincentNowak/PhD_thesis/blob/main/Chapter_2/Final_bigslice_query_sqlite_analyses_110223.ipynb). Using the Pfam-vectors created by BiG-SLICE for each BGC, the minimum cosine distance of all 2,670 BGCs relative to BGCs in the BiG-FAM database ^96^ inherent to BiG-SLICE was calculated. BGCs with a minimum cosine distance (min(d_BiG-FAM_)) larger than 0.2 were considered novel as previously determined.^23^ An all-to-all cosine distance was then also calculated between the 2,670 BGCs identified in this study and the MiBIG database.^97^ Gene Cluster Families (GCFs) and Gene Cluster Clans (GCCs) were constructed from this calculation with the AgglomerativeClustering function (average linkage) from the scikit-learn python library ^98^ and using the 0.2 and 0.8 cosine distance cutoffs for GCFs and GCCs respectively.^23^

## Acknowledgements

We thank Matt Storey and Manuel Blank for their help with the sponge sample collection and preparation. This work was supported by a Victoria University of Wellington Doctoral scholarship, the Health Research Council of New Zealand (contract 16/172), the Royal Society of New Zealand Te Apārangi (contract RDF-VUW1601) and the Ministry for Business Innovation and Employment (contracts RTVU1908 and UOAX2010). We would also like to thank the Centre for Academic Development at Victoria University of Wellington, in particular Andre Geldenhuis, for maintaining the Rāpoi high performance computing cluster, where computations for this work were performed.

