## supplementary information for "Microbial communities associated with marine sponges from diverse geographic locations harbour biosynthetic novelty"

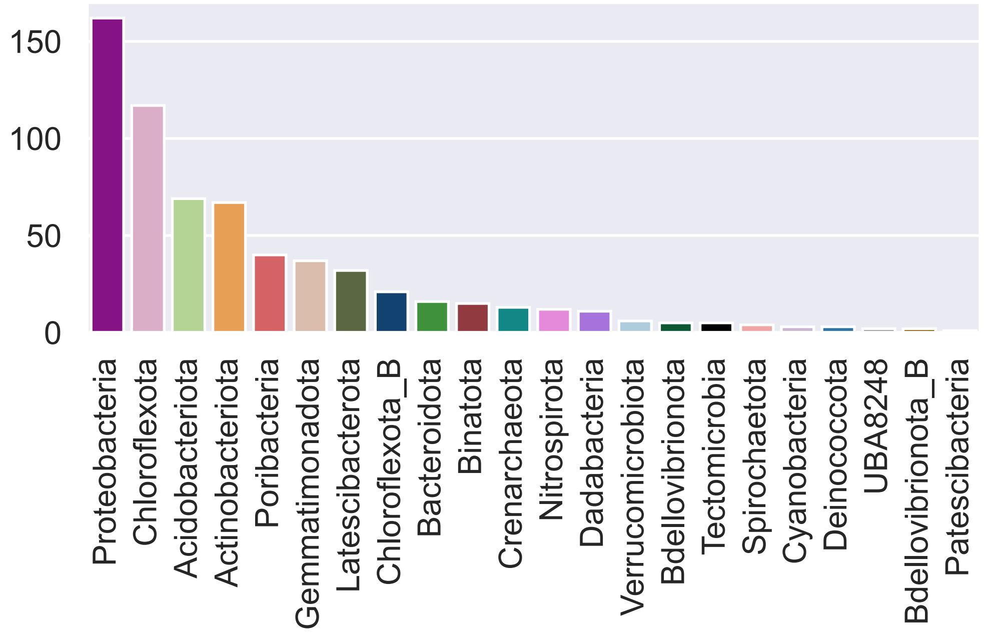


**Figure S1** Bar graph of the number of MAGs identified for each phylum across all samples

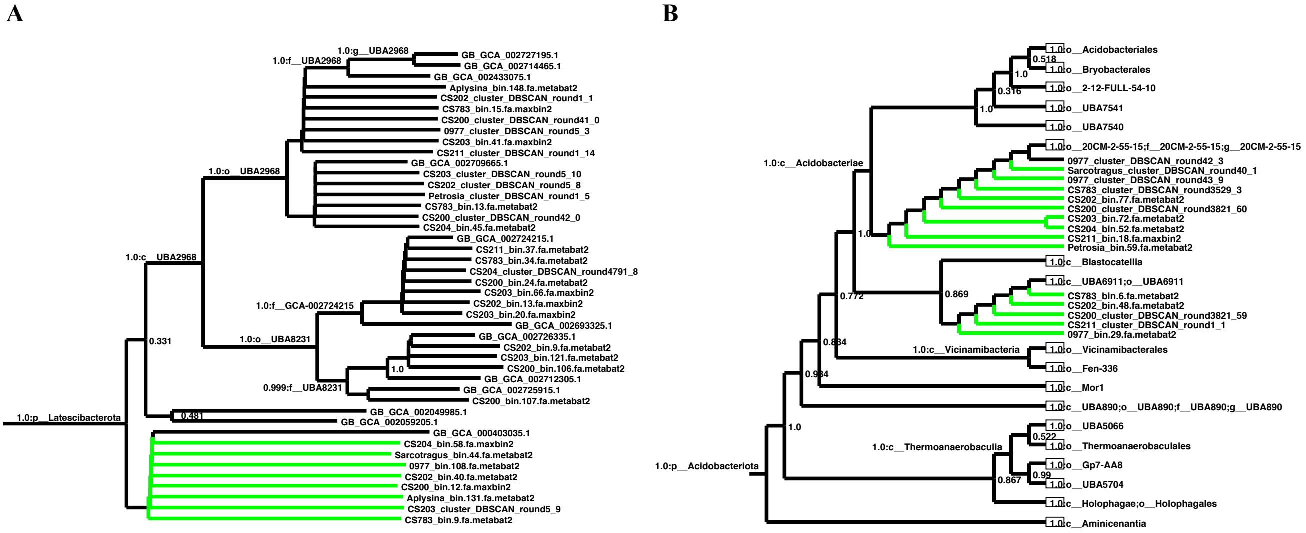


**Figure S2** Pruned phylogenetic tree extracted from the lager phylogenetic created by GTDB-Tk for all 643 MAGs. Nodes in green are MAGs identified in this study and highlight the novelty added in the phyla **A** Latescibacterota and **B** Acidobacteriota

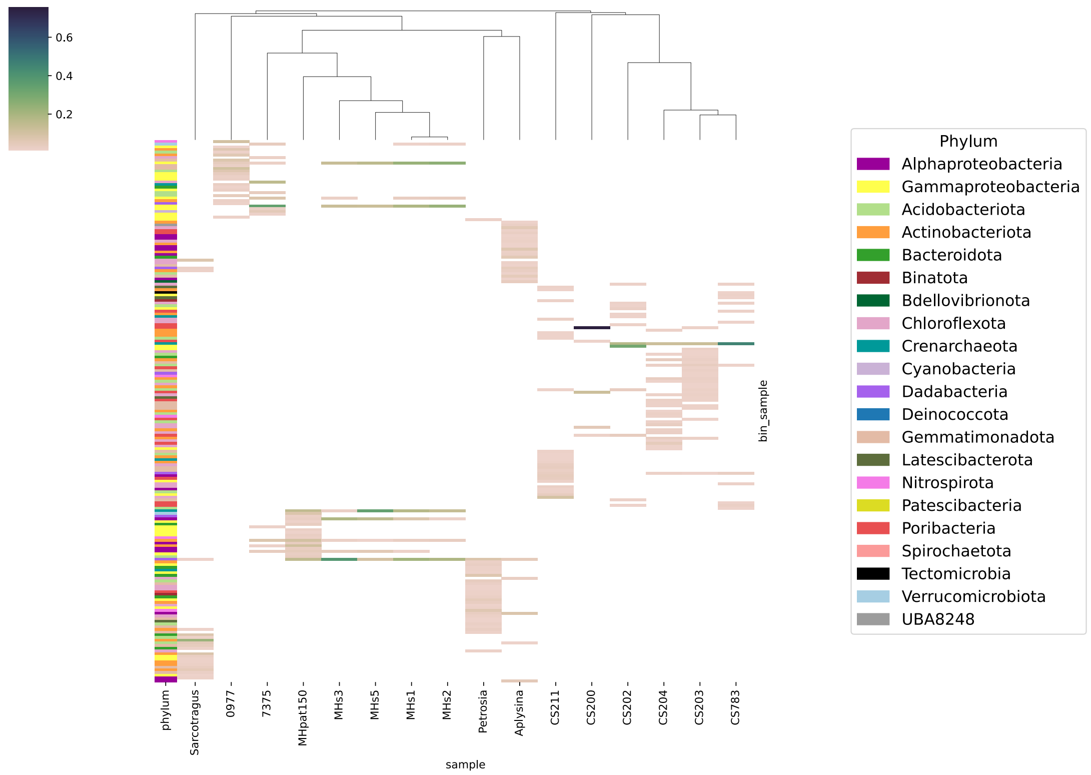


**Figure S3**: Abundance-based cosine clustering of all MAGs with a relative abundance of more than 1% in any of the samples

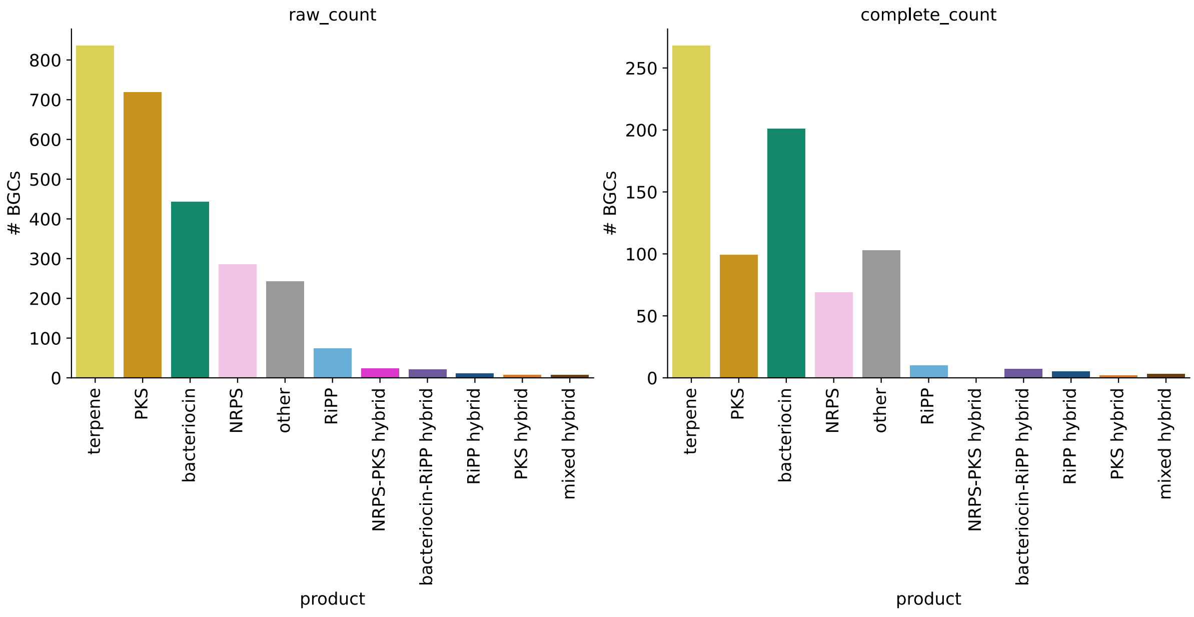


**Figure S4**: Number of BGCs identified across all 16 samples. Raw total on left and complete BGCs only on right. BGC classes/product types are summarised as outlined in Table S5

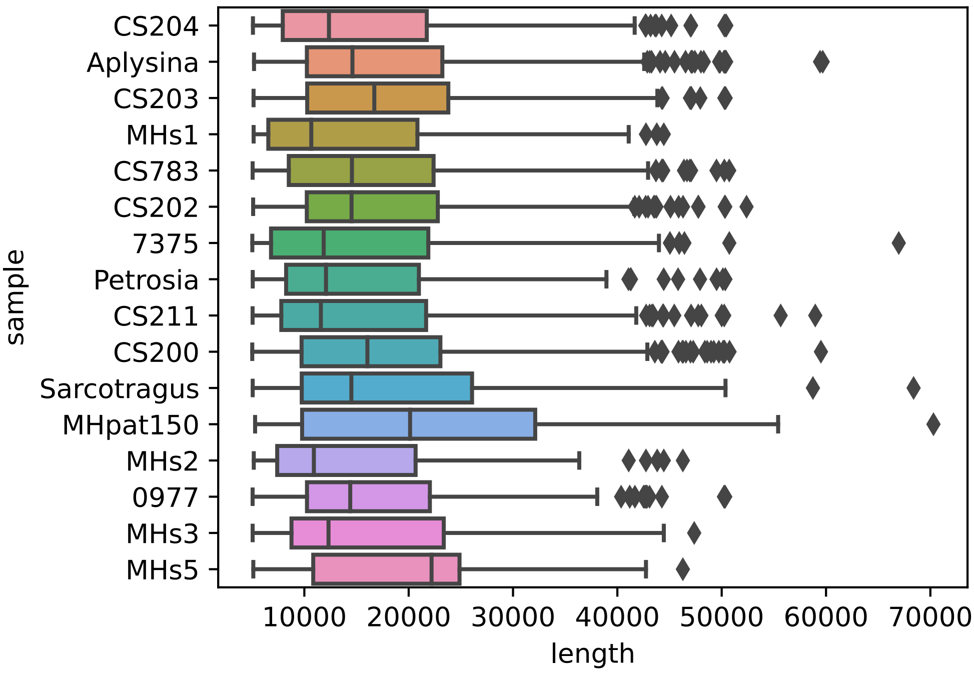


**Figure S5**: Boxplot of BGC lengths (*n* = 2,670) by sample

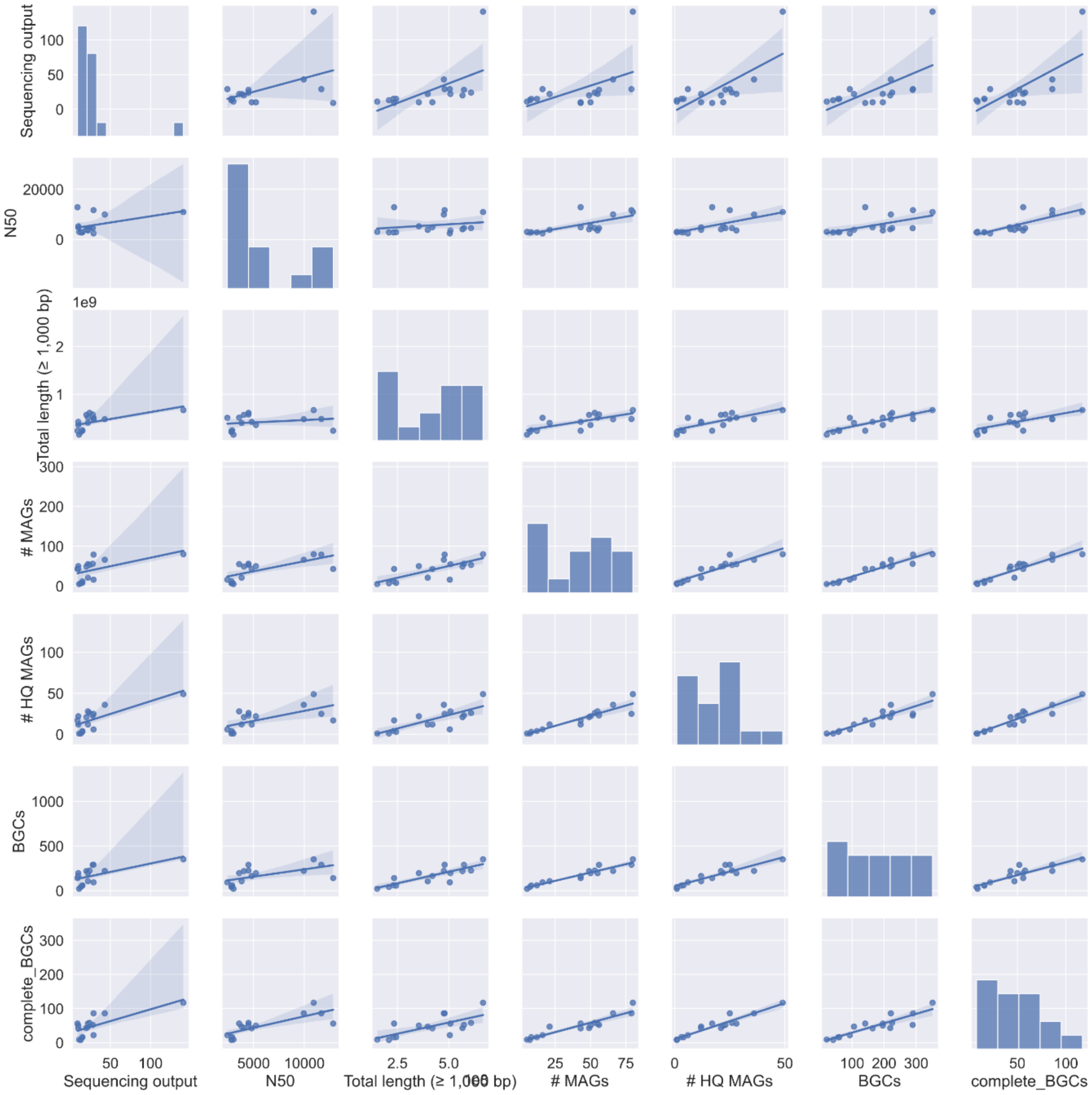


**Figure S6**: Pair plot relating general summary statistics to each other in a pairwise fashion with 95% confidence interval displayed as a blue shaded area. Raw data is visible in Table A9. Note that Aplysina is technically an outlier in terms of the sequencing output but was included here for completeness and since trends are unchanged if excluded. Sequencing output does not correlate well with any of the final result metrics. Total sequencing length correlates better than N50 with the downstream result metrics (number of MAGs, HQ MAGs, all BGCs and complete BGCs), putatively because it gives an indication of the size of the metagenome sequenced.

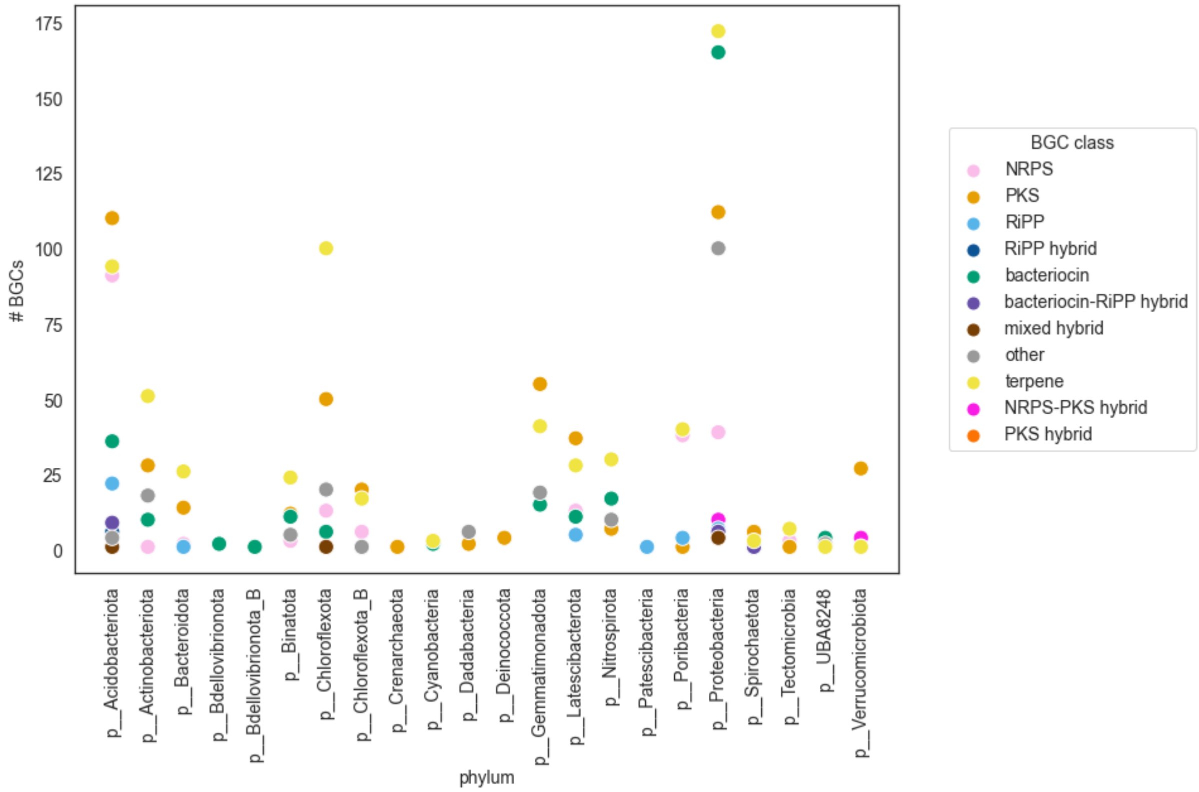


**Figure S7**: BGCs by phylum identified across all 16 samples

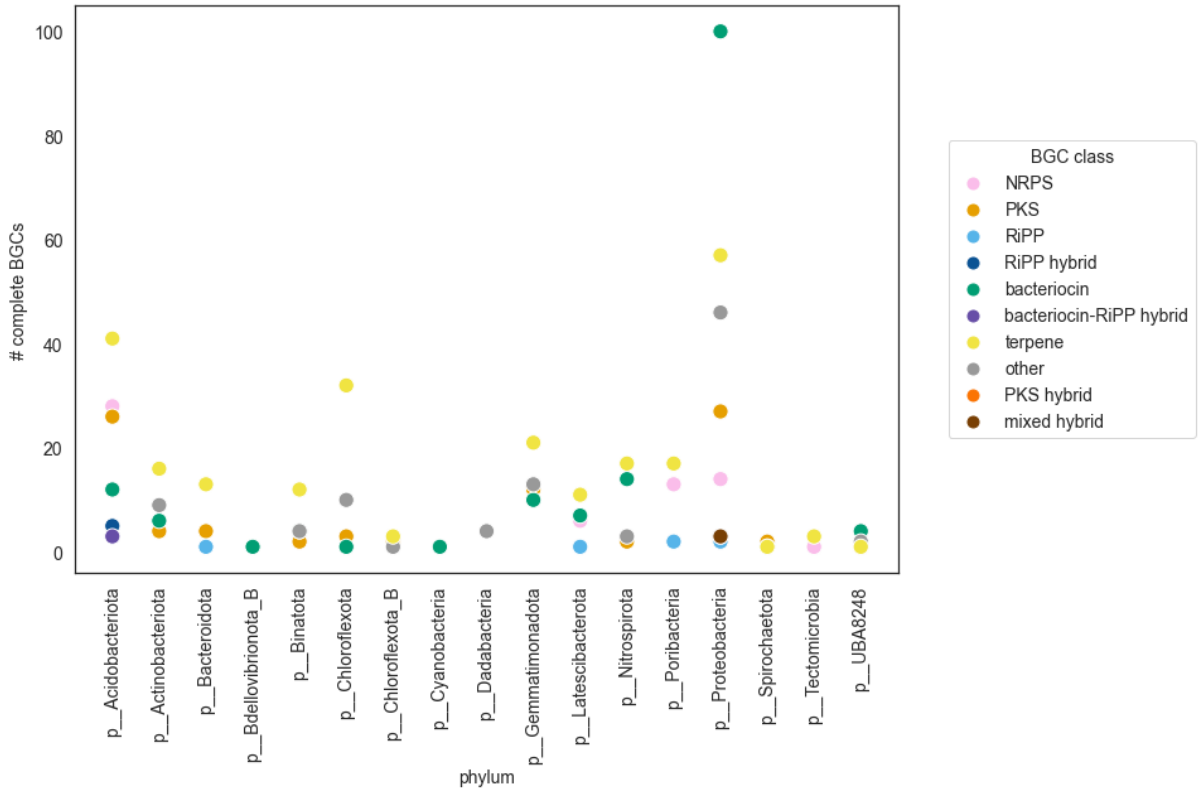


**Figure S8**: Complete BGCs (not on a contig edge) by phylum across all 16 samples

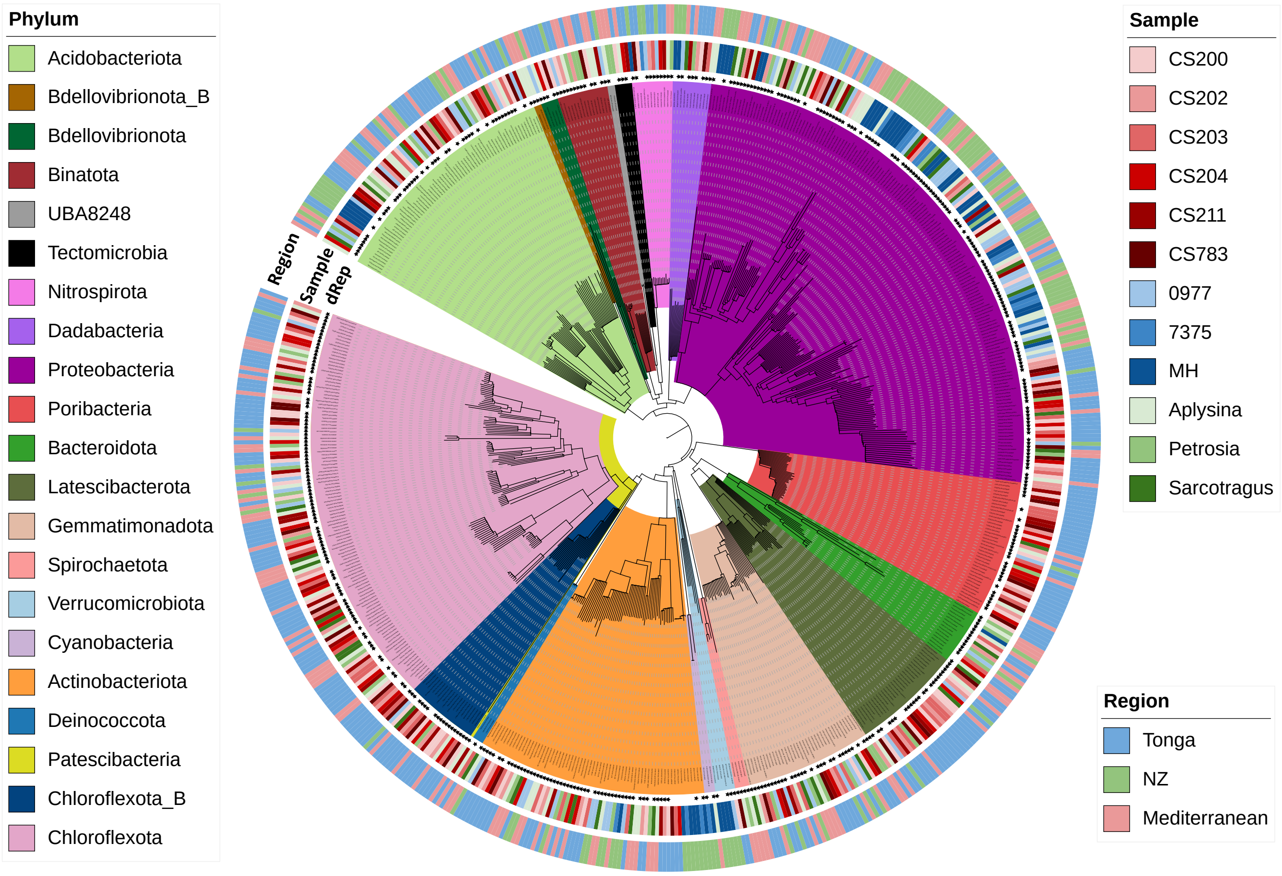


**Figure S9** Phylogenetic tree showing the 643 MAGs identified from the 16 marine sponge metagenomes. Black stars visible in the 'dRep' ring indicate MAGs that were identified as unique 'species' after dereplication of all 643 MAGs. The sample and region rings indicate the sample the MAG originated from and the region the sponge originated from respectively. Orientation of the phylogenetic tree is identical to Figure 14A.

A

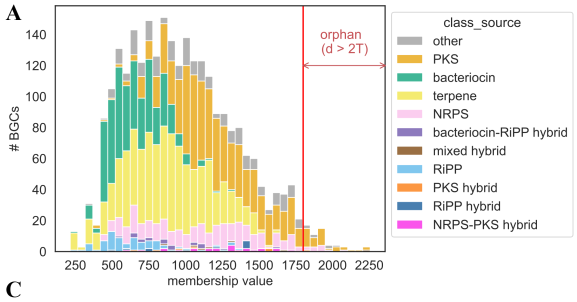


B

| product | count | Fraction fragmented |
|---|---|---|
| PKS | 39 | 0.28 |
| NRPS | 11 | 0.18 |
| other | 7 | 0.14 |
| Mixed hybrid | 1 | 0 |
| PKS hybrid | 1 | 1 |

C D

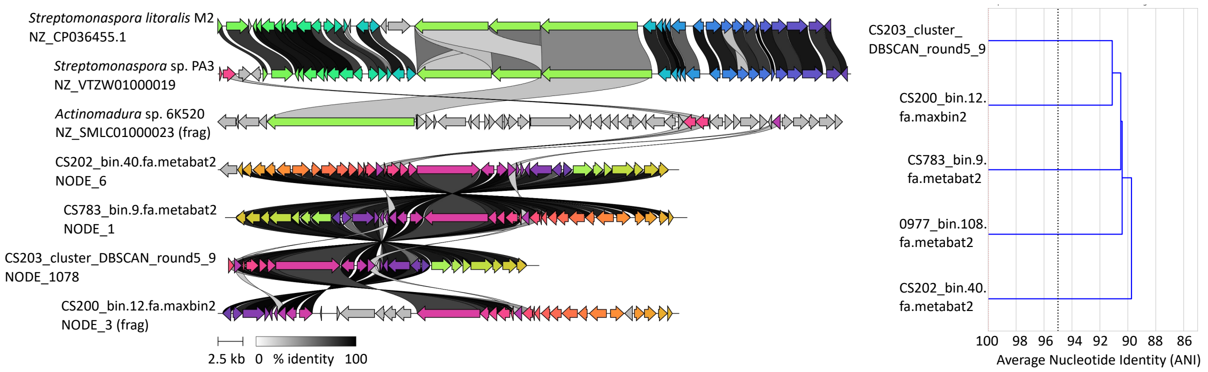


**Figure S10** **A** Histogram of BiG-SLICE membership values of all 2,670 BGCs **B** Number of orphan BGCs by product class **C** GCF_08405 visualised with clinker to show gene similarity between the three BGCs comprising the original GCF (labelled with species identified from) and the four BGCs from this work attribute to the GCF where fragmented BGCs are indicated as (frag) **D** Primary cluster produced by dRep containing the four 'species-level' MAGs the previously mentioned four BGCs were associated with. All MAGs in the cluster could not be classified below the Latescibacteriota phylum level.

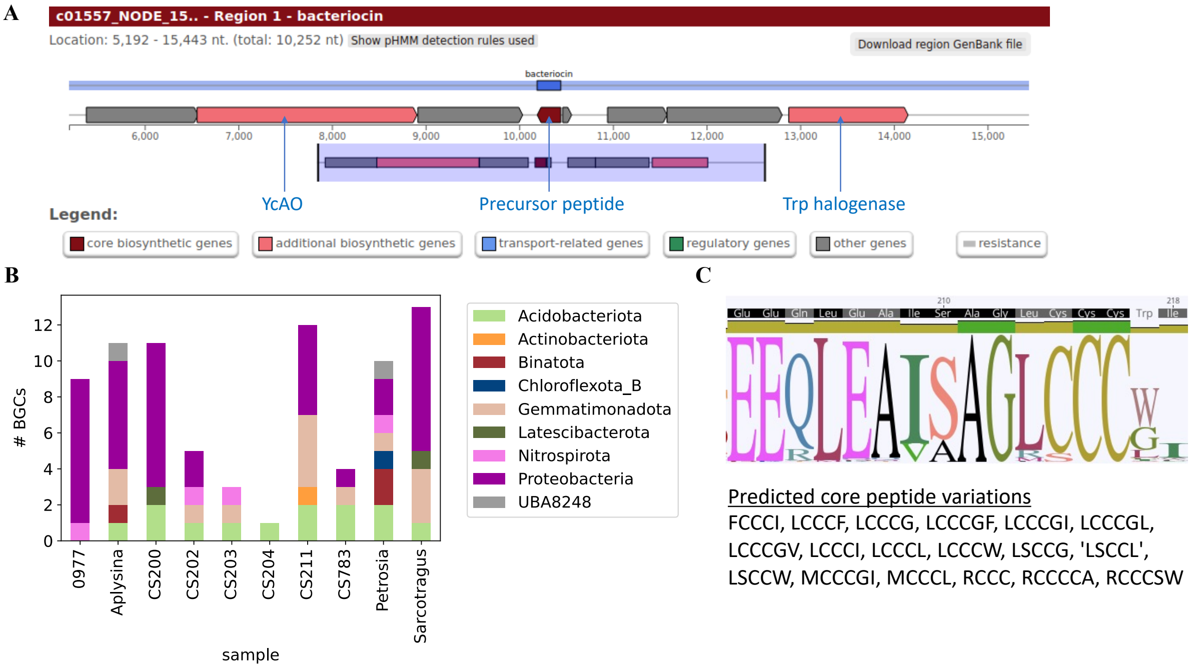


**Figure S11** Sponge-derived RiPP proteusins previously identified (Nguyen et al. 2021) where **A** is an example BGC, **B** shows the number of BGCs identified per sample and bacterial phylum and **C** shows the conserved peptide motif including some newly identified variants.

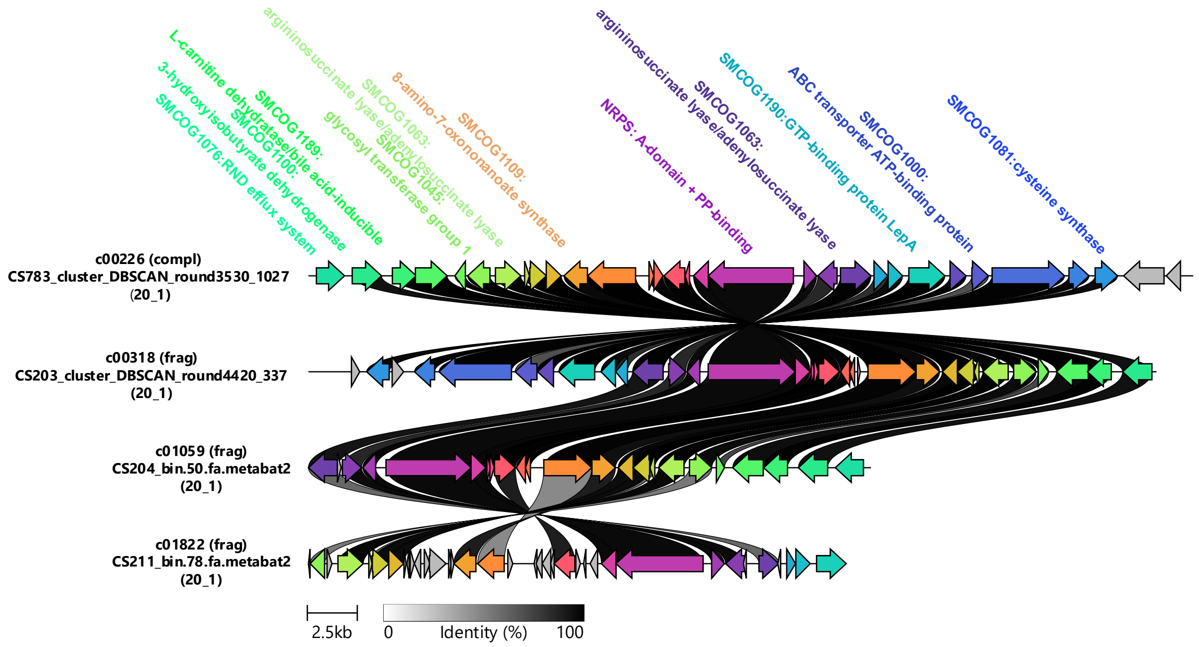


**Figure S12** GCF_18 associated with a single dRep species cluster (20_1) present in four of the six Tongan sponges

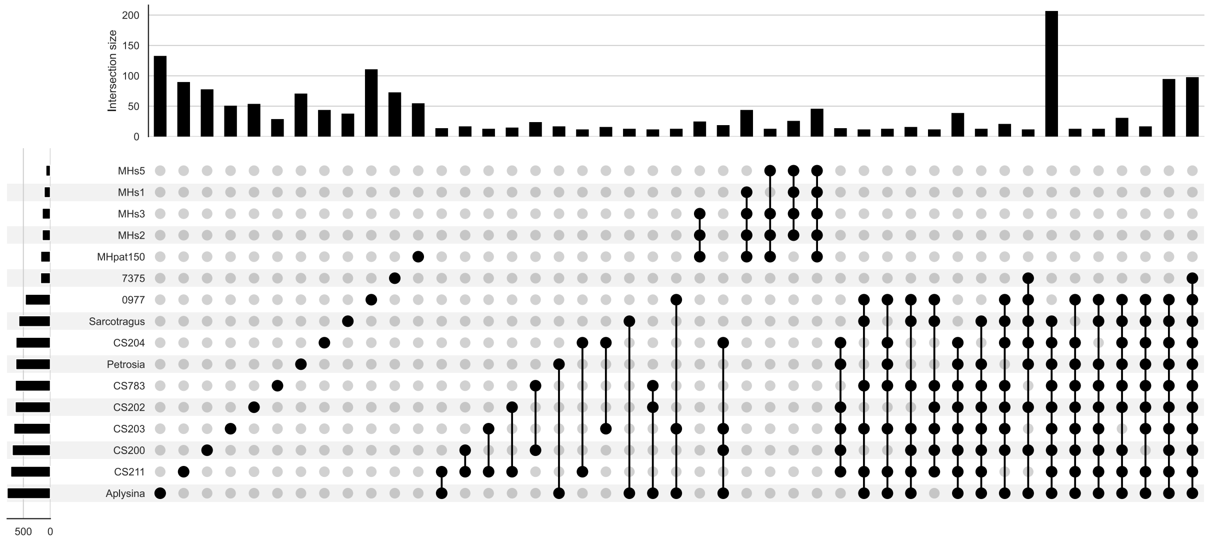


**Figure S13** Upset plot of GCFs shared between any combination of samples. Only values above 10 shared GCFs are included in this graph.

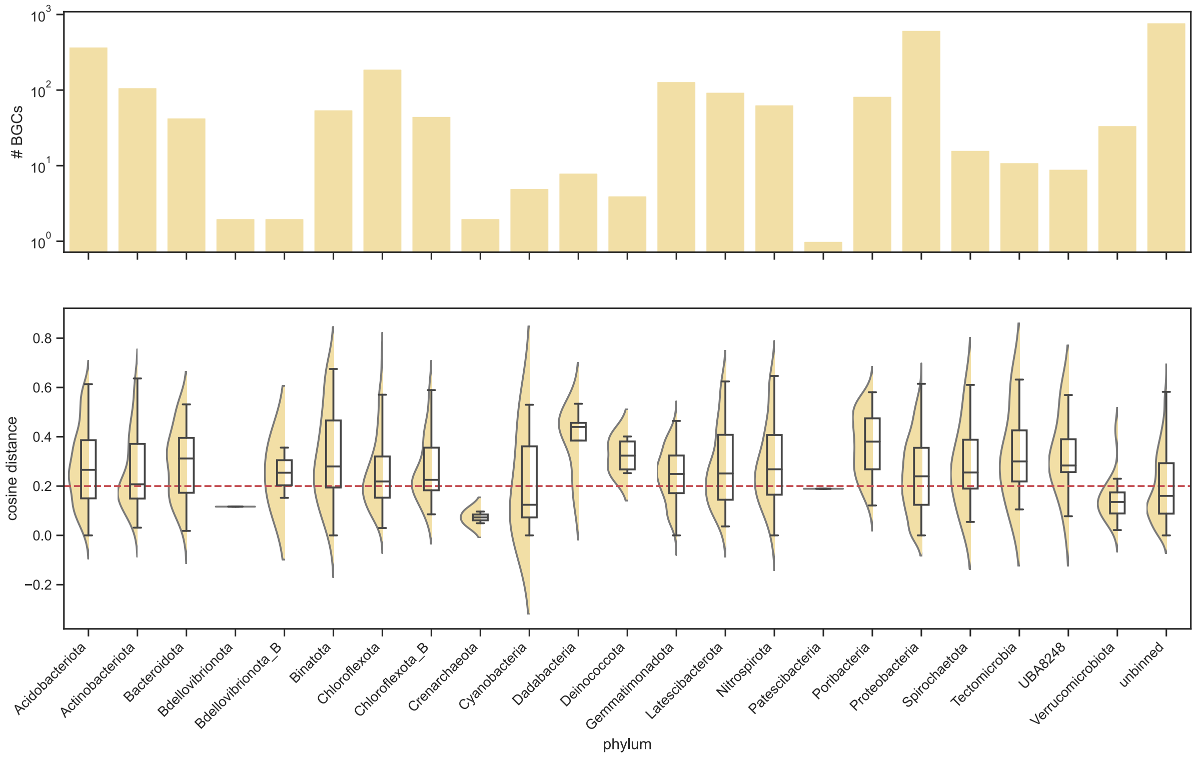


**Figure S14** min($d_{BiG-FAM}$) by bacterial phylum with number of BGCs (log scale) in top panel and boxplot of median and quartiles as well as min($d_{BiG-FAM}$) value distribution (relative to number of BGCs in respective phylum) in bottom panel.

| Sample | Origin and collection date | Date sequenced | Sequencing Technology | Sequencing library | Reads returned | Gigabases returned |
|---|---|---|---|---|---|---|
| *Cacospongia mycofijiensis* (CS783) | Cathedral Cave, 'Eua, Tonga on 07/06/2016 | 11/09/2017 | Illumina HiSeq 4000 (Annorad) | 2×150bp paired- end (TruSeq, PCR free) | 70,524,102 | 10.58 |
| CS200 (834) | Pete's cave, 'Eua, Tonga on 08/06/2016 | 01/05/2018 | Illumina HiSeq 4000 (Annorad) | 2×150bp paired- end (TruSeq, PCR free) | 194,945,500 | 29.24 |
| CS202 (837) | Pete's cave, 'Eua, Tonga on 08/06/2016 | 01/05/2018 | Illumina HiSeq 4000 (Annorad) | 2×150bp paired- end (TruSeq, PCR free) | 149,578,212 | 22.44 |
| CS203 (839) | Pete's cave, 'Eua, Tonga on 08/06/2016 | 01/05/2018 | Illumina HiSeq 4000 (Annorad) | 2×150bp paired- end (TruSeq, PCR free) | 163,853,646 | 24.58 |
| CS204 (841) | Pete's cave, 'Eua, Tonga on 08/06/2016 | 01/05/2018 | Illumina HiSeq 4000 (Annorad) | 2×150bp paired- end (TruSeq, PCR free) | 136,937,548 | 20.54 |
| CS211 (854) | Pete's cave, 'Eua, Tonga on 09/06/2016 | 01/05/2018 | Illumina HiSeq 4000 (Annorad) | 2×150bp paired- end (TruSeq, PCR free) | 186,895,268 | 28.04 |
| MHPat150 (Storey et al. 2020) | Capsize Point, Malborough Sounds, New Zealand Nov 2014 | Aug 2018 | Illumina HiSeq 4000 (Genewiz) | 2×150bp paired- end (TruSeq) | 147,632,620 | 22.14 |
| MHs1 (Storey et al. 2020) | Pelorus Sound, Malborough Sounds, New Zeland May 2003 | Aug 2018 | Illumina HiSeq 4000 (Genewiz) | 2×150bp paired- end (TruSeq) | 90,573,844 | 13.59 |
| MHs2 (Storey et al. 2020) | Pelorus Sound, Malborough Sounds, New Zeland May 2003 | Aug 2018 | Illumina HiSeq 4000 (Genewiz) | 2×150bp paired- end (TruSeq) | 102,500,088 | 15.38 |
| MHs3 (Storey et al. 2020) | Pelorus Sound, Malborough Sounds, New Zeland May 2003 | Aug 2018 | Illumina HiSeq 4000 (Genewiz) | 2×150bp paired- end (TruSeq) | 101,083,926 | 15.16 |
| MHs5 (Storey et al. 2020) | Pelorus Sound, Malborough Sounds, New Zeland May 2003 | Aug 2018 | Illumina HiSeq 4000 (Genewiz) | 2×150bp paired- end (TruSeq) | 76,506,312 | 11.48 |

| | | | | | | |
|---|---|---|---|---|---|---|
| MNP0977 | Stephens Island, New Zealand (S40.6583333° E174°) | 16/10/2019 | Illumina HiSeq 4000 (Genewiz) | 2×150bp paired- end (TruSeq) | 289,571,846 | 43.44 |
| MNP7375 | Keneparu Sound, New Zealand (S41.2193167° E173.8717333°) 24/05/2004 | 16/10/2019 | Illumina HiSeq 4000 (Genewiz) | 2×150bp paired- end (TruSeq) | 199,352,714 | 29.90 |
| *Aplysina aerophoba* (Horn et al. 2016) | Piran, Slovenia 07/05/2013 | 2013? | Illumina HiSeq 2000 (DOE JGI) | 2×150bp paired- end library type unknown | 945,906,728 | 141.89 |
| *Petrosia ficiformis* (Horn et al. 2016) | Milos, Greece (N36.76759° E24.51422°) 25/05/2013 | 2013? | Illumina MiSeq Personal Sequencer (GATC Biotech AG) | 2×250bp paired- end (FastQC says 251) library type unknown | 41,383,564 | 10.35 |
| *Sarcotragus foetidus* (Horn et al. 2016) | Milos, Greece (N36.76759° E24.51422°) 25/05/2013 | 2013? | Illumina MiSeq Personal Sequencer (GATC Biotech AG) | 2×300bp paired- end (FastQC says 301) library type unknown | 32,672,426 | 9.80 |

**Table S1** Sample and sequencing metadata table, all sponges collected by SCUBA divers. Sequencing output for all samples was standardised to Gigabases (Gb) rather than Gigabase pairs (Gbp), where Gb = (forward reads + reverse reads) * sequence length.

| **Sponge** | **Historical taxonomy** | **18S rRNA taxonomy** | **Align. Qual.** | **Based on complete gene** | **Nearest SILVA alignment reference (alignment quality)** | **Taxonomy of nearest SILVA alignment** |
|---|---|---|---|---|---|---|
| MNP0977 | Keratosa, Dictyoceratida, Irciniidae, *Psammocinia* "unknown spp" | Keratosa, Dictyoceratida | 99 | Yes (1802 bp) | AY734448.1.1.1791 (0.982) | Keratosa, Dictyoceratida, Irciniidae, *Ircinia felix* |
| MNP7375 | Heteroscleromorpha, Halichondrida, Halichondriidae, *Halichondria* "n. sp. 4" | n/a | 99 | No (1356 bp) | KC902238.1.1.1744 (0.987) | Heteroscleromorpha, Halichondrida, Halichondriidae, *Halichondria panicea* |
| MHPat | Heteroscleromorpha, Poecilosclerida, Mycalidae, *Mycale hentscheli* | Poecilosclerida | 99 | Yes (1817 bp) | AY737643.1.1.1803 (0.979) | Heteroscleromorpha, Poecilosclerida, Mycalidae, *Mycale sp. sp16* |
| MHs1 | Heteroscleromorpha, Poecilosclerida, Mycalidae, *Mycale hentscheli* | Poecilosclerida | 99 | No (1617 bp) | AY737643.1.1.1803 (0.987) | Heteroscleromorpha, Poecilosclerida, Mycalidae, *Mycale sp. sp16* |
| MHs2 | Heteroscleromorpha, Poecilosclerida, Mycalidae, *Mycale hentscheli* | Poecilosclerida | 99 | No (1237 bp) | AY737643.1.1.1803 (0.985) | Heteroscleromorpha, Poecilosclerida, Mycalidae, *Mycale sp. sp16* |
| MHs3 | Heteroscleromorpha, Poecilosclerida, Mycalidae, *Mycale hentscheli* | Poecilosclerida | 99 | Yes (1817 bp) | AY737643.1.1.1803 (0.979) | Heteroscleromorpha, Poecilosclerida, Mycalidae, *Mycale sp. sp16* |
| MHs5 | Heteroscleromorpha, Poecilosclerida, Mycalidae, *Mycale hentscheli* | Poecilosclerida | 97 | No (513 bp) | KX622151.1.1.1767 (0.926) | Heteroscleromorpha, Poecilosclerida, Mycalidae, *Mycale sanguinea* |
| CS783 | Keratosa, Dictyoceratida, Thorectidae, *Cacospongia mycofijiensis* | Dictyoceratida | 99 | Yes (1802 bp) | AY34448.1.1.1791 (0.976) | Keratosa, Dictyoceratida, Irciniidae, *Ircinia felix* |
| CS200 | n/a | Dictyoceratida | 99 | Yes (1766 bp) | AY34448.1.1.1791 (0.981) | Keratosa, Dictyoceratida, Irciniidae, *Ircinia felix* |
| CS202 | n/a | Dictyoceratida | 99 | Yes (1805 bp) | KX894466.1.1.1801 (0.974) | Keratosa, Dictyoceratida, Thorectidae, |

| | | | | | | |
|---|---|---|---|---|---|---|
| | | | | | | *Dactylospongia sp. DAS.2* |
| CS203 | n/a | Verongiida | 99 | Yes (1798 bp) | AY591801.1.1.1799 (0.993) | Veronigmorpha, Verongiida, Aplysinidae, *Aplysina archeri* |
| CS204 | n/a | Verongiida | 99 | Yes (1798 bp) | AY591801.1.1.1799 (0.993) | Veronigmorpha, Verongiida, Aplysinidae, *Aplysina archeri* |
| CS211 | n/a | Dictyoceratida | 99 | No (1208 bp) | AY734448.1.1.1791 (0.988) | Keratosa, Dictyoceratida, Irciniidae, *Ircinia felix* |
| Aplysina | Veronigmorpha, Verongiida, Aplysinidae, *Aplysina aerophoba* | Verongiida | 99 | Yes (1798 bp) | AY591801.1.1.1799 (0.993) | Veronigmorpha, Verongiida, Aplysinidae, *Aplysina archeri* |
| Petrosia | Heteroscleromorpha, Haplosclerida, Petrosiidae, *Petrosia ficiformis* | Haplosclerida | 99 | Yes (2023 bp) | KX622161.1.1.1975 (0.969) | Heteroscleromorpha, Haplosclerida, Petrosiidae, *Petrosia ficiformis* |
| Sarcotragus | Keratosa, Dictyoceratida, Irciniidae, *Sarcotragus foetidus* | Dictyoceratida | 99 | No (911 bp) | AY734448.1.1.1791 (0.977) | Keratosa, Dictyoceratida, Irciniidae, *Ircinia felix* |

**Table S2** MNP0977 and MNP7375 identified by NIWA, MH samples identified by?, CS783 identified by? 18S rRNA sequences extracted using barrnap v.0.9 and SINA commandline tool v.1.6.1 using the SILVA_138_SSURef_NR99. All sponges identified as Demospongiae.

| Sample | Sequencing output | N50 (bp) | Total length (≥ 1,000 bp) | # MAGs | # HQ MAGs | BGCs | complete_BGCs |
|---|---|---|---|---|---|---|---|
| **MNP0977** | 43.44 | 9948 | 475953997 | 66 | 36 | 221 | 86 |
| **MNP7375** | 29.9 | 2409 | 504957411 | 16 | 6 | 93 | 22 |
| **MHpat150** | 22.1 | 3816 | 397458873 | 21 | 12 | 106 | 47 |
| **MHs1** | 13.59 | 2807 | 208520661 | 7 | 1 | 41 | 8 |
| **MHs2** | 15.38 | 2889 | 243615086 | 8 | 3 | 57 | 16 |
| **MHs3** | 15.16 | 2819 | 230585775 | 12 | 4 | 60 | 16 |
| **MHs5** | 11.48 | 3011 | 151400149 | 5 | 1 | 21 | 9 |
| **CS200** | 29.24 | 11682 | 479674948 | 79 | 25 | 290 | 86 |
| **CS202** | 22.44 | 3575 | 507780759 | 55 | 28 | 196 | 56 |
| **CS203** | 24.58 | 4498 | 608488494 | 53 | 26 | 225 | 58 |
| **CS204** | 20.54 | 4044 | 566726134 | 49 | 21 | 219 | 43 |
| **CS211** | 28.04 | 4491 | 573988474 | 56 | 23 | 289 | 52 |
| **CS783** | 10.58 | 4814 | 420034886 | 43 | 12 | 164 | 42 |
| ***A. aerophoba*** | 141.89 | 10926 | 666674772 | 80 | 49 | 351 | 117 |
| ***P. ficiformis*** | 10.35 | 5227 | 354237708 | 50 | 22 | 196 | 50 |
| ***S. foetidus*** | 9.8 | 12850 | 233760078 | 43 | 17 | 141 | 56 |

**Table S3:** Raw summary statistics for results from the MetaSing pipeline. Assembly statistics for all assemblies as determined by Quast v.5.0.2 [348] run with the -m 1000 flag to base statistics on contigs >1000 bp. The assemblies originally reported for *A. aerophoba*, *P. ficiformis* and *S. foetidus* produced an N50 and a total length (≥ 1000 bp) of 8,958 and 489,999,481, 3,381 and 226,772,563 and 9,706 and 190,159,175 respectively. The assemblies reported for Mhs1 (ASM1226765v1), MHs2 (ASM1226770v1), MHs3 (ASM1226336v1) and MHs5 (ASM1226332v1) produced comparable statistics but note that statistics reported on NCBI are based on all contig sizes.

| Phylum | Collective 643 MAGs | Dereplicated 510 MAGs | Percentage change (-%) |
|---|---|---|---|
| Proteobacteria | 162 | 134 | 17.3 |
| Chloroflexota | 117 | 98 | 16.2 |
| Acidobacteriota | 69 | 45 | 34.8 |
| Actinobacteriota | 67 | 54 | 19.4 |
| Poribacteria | 40 | 29 | 27.5 |
| Gemmatimonadota | 37 | 30 | 18.9 |
| Latescibacterota | 32 | 24 | 25.0 |
| Chloroflexota_B | 21 | 18 | 14.3 |

| | | | |
|---|---|---|---|
| Bacteroidota | 16 | 14 | 12.5 |
| Binatota | 15 | 13 | 13.3 |
| Crenarchaeota | 13 | 11 | 15.4 |
| Nitrospirota | 12 | 10 | 16.7 |
| Dadabacteria | 11 | 8 | 27.2 |
| Bdellovibrionota | 5 | 4 | 20.0 |
| Verrucomicrobiota | 6 | 4 | 33.3 |
| Tectomicrobia | 5 | 3 | 40.0 |
| Spirochaetota | 4 | 4 | 0.0 |
| Cyanobacteria | 3 | 2 | 33.3 |
| Deinococcota | 3 | 1 | 66.7 |
| UBA8248 | 2 | 1 | 50.0 |
| Bdellovibrionota_B | 2 | 2 | 0 |
| Patescibacteria | 1 | 1 | 1 |

**Table S4** Count of MAGs by phylum in the collective set of 643 MAGs identified from all 16 samples and in the dereplicated set of 510 species-level MAGs

| Product | Raw count | Complete count | Summarised as |
|---|---|---|---|
| terpene | 836 | 268 | terpene |
| T1PKS | 538 | 75 | PKS |
| bacteriocin | 443 | 201 | bacteriocin |
| NRPS-like | 248 | 63 | NRPS |
| betalactone | 123 | 59 | other |
| T3PKS | 76 | 12 | PKS |
| transAT-PKS-like | 53 | 0 | PKS |
| arylpolyene | 42 | 9 | PKS |
| NRPS | 36 | 6 | NRPS |
| ectoine | 36 | 16 | other |
| phosphonate | 35 | 9 | other |
| lassopeptide | 31 | 3 | RiPP |
| lanthipeptide | 19 | 4 | RiPP |
| other | 18 | 7 | other |
| LAP | 17 | 2 | RiPP |
| hserlactone | 14 | 6 | other |
| NRPS-transAT-PKS-like | 8 | 0 | NRPS-PKS hybrid |
| NRPS-T1PKS | 8 | 0 | NRPS-PKS hybrid |
| bacteriocin-proteusin | 7 | 3 | bacteriocin-RiPP hybrid |
| nucleoside | 7 | 0 | other |
| bacteriocin-lanthipeptide | 6 | 2 | bacteriocin-RiPP hybrid |
| TfuA-related | 5 | 1 | RiPP |
| LAP-lassopeptide | 5 | 5 | RiPP hybrid |
| LAP-bacteriocin | 5 | 1 | bacteriocin-RiPP hybrid |
| oligosaccharide | 5 | 5 | other |
| bacteriocin-terpene | 4 | 2 | mixed hybrid |
| NRPS-like-transAT-PKS-like | 4 | 0 | NRPS-PKS hybrid |
| PKS-like | 4 | 0 | PKS |
| LAP-thiopeptide | 3 | 0 | RiPP hybrid |
| resorcinol | 3 | 2 | other |
| NRPS-like-T1PKS | 3 | 0 | NRPS-PKS hybrid |
| siderophore | 3 | 1 | other |
| PKS-like-T3PKS | 2 | 0 | PKS hybrid |
| bacteriocin-thiopeptide | 2 | 0 | bacteriocin-RiPP hybrid |
| ladderane | 2 | 0 | other |
| LAP-proteusin | 2 | 0 | RiPP hybrid |
| T1PKS-hglE-KS | 2 | 1 | PKS hybrid |
| TfuA-related-bacteriocin-proteusin | 1 | 1 | bacteriocin-RiPP hybrid |
| transAT-PKS | 1 | 0 | PKS |
| NRPS-transAT-PKS | 1 | 0 | NRPS-PKS hybrid |
| phosphonate-terpene | 1 | 1 | mixed hybrid |
| proteusin | 1 | 0 | RiPP |
| hglE-KS | 1 | 1 | PKS |
| T2PKS | 1 | 0 | PKS |
| arylpolyene-resorcinol | 1 | 1 | PKS hybrid |
| NRPS-like-betalactone | 1 | 0 | mixed hybrid |
| TfuA-related-proteusin | 1 | 0 | RiPP hybrid |
| CDPS | 1 | 0 | NRPS |
| arylpolyene-ladderane | 1 | 0 | PKS hybrid |
| head_to_tail | 1 | 0 | RiPP |

| | | | |
|---|---|---|---|
| NRPS-like-terpene | 1 | 0 | mixed hybrid |
| T3PKS-transAT-PKS | 1 | 0 | PKS hybrid |

**Table S5** Number of BGCs identified across all 16 marine sponge metagenomes with the number of raw number of BGCs (complete and incomplete) and number of complete BGCs given per product class/BGC class as identified by antiSMASH5. Due to the large number of classes and to allow easier visualisation, a summary term was created.

| | # MAGs | # BGCs | # complete BGCs | Avg. BGCs per MAG | Avg. complete BGCs per MAG |
|---|---|---|---|---|---|
| p__Verrucomicrobiota | 6 | 34 | 0 | 5.67 | 0 |
| p__Acidobacteriota | 69 | 373 | 118 | 5.41 | 1.71 |
| p__Nitrospirota | 12 | 64 | 36 | 5.33 | 3 |
| p__UBA8248 | 2 | 9 | 8 | 4.5 | 4 |
| p__Spirochaetota | 4 | 16 | 5 | 4 | 1.25 |
| p__Proteobacteria | 162 | 619 | 254 | 3.82 | 1.57 |
| p__Binatota | 15 | 55 | 24 | 3.67 | 1.6 |
| p__Gemmatimonadota | 37 | 130 | 56 | 3.51 | 1.51 |
| p__Latescibacterota | 32 | 94 | 36 | 2.94 | 1.12 |
| p__Bacteroidota | 16 | 43 | 18 | 2.69 | 1.12 |
| p__Tectomicrobia | 5 | 11 | 4 | 2.2 | 0.8 |
| p__Chloroflexota_B | 21 | 45 | 8 | 2.14 | 0.38 |
| p__Poribacteria | 40 | 83 | 32 | 2.08 | 0.8 |
| p__Cyanobacteria | 3 | 5 | 1 | 1.67 | 0.33 |
| p__Chloroflexota | 117 | 190 | 47 | 1.62 | 0.4 |
| p__Actinobacteriota | 67 | 108 | 35 | 1.61 | 0.52 |
| p__Deinococcota | 3 | 4 | 0 | 1.33 | 0 |
| p__Bdellovibrionota_B | 2 | 2 | 1 | 1 | 0.5 |
| p__Patescibacteria | 1 | 1 | 0 | 1 | 0 |
| p__Dadabacteria | 11 | 8 | 4 | 0.73 | 0.36 |
| p__Bdellovibrionota | 5 | 2 | 0 | 0.4 | 0 |
| p__Crenarchaeota | 13 | 2 | 0 | 0.15 | 0 |

**Table S6** Number of BGCs found per MAGs at the phylum level (as identified by GTDB-Tk). Note that MAG counts are based on all 643 MAGs to account for MAGs that do not have BGCs associated with them.

| **Phylum** | **BGC % mean** |
|---|---|
| Verrucomicrobiota | 4.78 |
| Nitrospirota | 4.40 |
| Acidobacteriota | 2.51 |
| Acidobacteriota | 2.33 |
| UBA8248 | 2.33 |
| Dadabacteria | 2.32 |
| Gemmatimonadota | 2.24 |
| Bacteroidota | 2.19 |
| Bdellovibrionota_B | 2.04 |
| Binatota | 1.95 |
| Proteobacteria | 1.81 |
| Chloroflexota_B | 1.69 |

| | |
|---|---|
| Spirochaetota | 1.63 |
| Latescibacterota | 1.55 |
| Chloroflexota | 1.34 |
| Cyanobacteria | 1.29 |
| Tectomicrobia | 1.25 |
| Deinococcota | 1.16 |
| Actinobacteriota | 1.15 |
| Poribacteria | 0.95 |
| Crenarchaeota | 0.68 |
| Bdellovibrionota | 0.38 |
| Patescibacteria | 0.36 |

**Table S7** Phylum mean of BGC % per MAG relative to genome size

| BGC class | # BGCs min($d_{BiG-FAM}$) ≤ 0.2 | # BGCs min($d_{BiG-FAM}$) > 0.2 | # BGCs | Novelty rate | Novelty rank | # BGCs rank |
|---|---|---|---|---|---|---|
| RiPP hybrid | 1 | 10 | 11 | 0.90909091 | 1 | 9 |
| mixed hybrid | 1 | 6 | 7 | 0.85714286 | 2 | 11 |
| other | 41 | 202 | 243 | 0.83127572 | 3 | 5 |
| bacteriocin-RiPP hybrid | 4 | 17 | 21 | 0.80952381 | 4 | 8 |
| NRPS | 68 | 217 | 285 | 0.76140351 | 5 | 4 |
| terpene | 244 | 592 | 836 | 0.70813397 | 6 | 1 |
| bacteriocin | 207 | 236 | 443 | 0.53273138 | 7 | 3 |
| RiPP | 40 | 34 | 74 | 0.45945946 | 8 | 6 |
| PKS | 534 | 185 | 719 | 0.25730181 | 9 | 2 |
| PKS hybrid | 6 | 1 | 7 | 0.14285714 | 10 | 11 |
| NRPS-PKS hybrid | 23 | 1 | 24 | 0.04166667 | 11 | 7 |

**Table S8** Rate of novel BGCs by BGC class as determined by min($d_{BiG-FAM}$)

| Phylum | # BGCs min($d_{BiG-FAM}$) ≤ 0.2 | # BGCs min($d_{BiG-FAM}$) > 0.2 | # BGCs | Novelty rate | Novelty rank | # BGCs rank |
|---|---|---|---|---|---|---|
| Deinococcota | 0 | 4 | 4 | 1 | 1 | 19 |
| Poribacteria | 8 | 75 | 83 | 0.90361446 | 2 | 8 |
| UBA8248 | 1 | 8 | 9 | 0.88888889 | 3 | 16 |
| Dadabacteria | 1 | 7 | 8 | 0.875 | 4 | 17 |
| Spirochaetota | 4 | 12 | 16 | 0.75 | 5 | 14 |
| Tectomicrobia | 3 | 8 | 11 | 0.72727273 | 6 | 15 |
| Chloroflexota_B | 14 | 31 | 45 | 0.68888889 | 7 | 11 |
| Acidobacteriota | 119 | 254 | 373 | 0.68096515 | 8 | 3 |
| Binatota | 18 | 37 | 55 | 0.67272727 | 9 | 10 |
| Nitrospirota | 21 | 43 | 64 | 0.671875 | 10 | 9 |
| Gemmatimonadota | 43 | 87 | 130 | 0.66923077 | 11 | 5 |
| Bacteroidota | 15 | 28 | 43 | 0.65116279 | 12 | 12 |
| Latescibacterota | 34 | 60 | 94 | 0.63829787 | 13 | 7 |
| Chloroflexota | 81 | 109 | 190 | 0.57368421 | 14 | 4 |

| | | | | | | |
|---|---|---|---|---|---|---|
| Proteobacteria | 267 | 352 | 619 | 0.56865913 | 15 | 2 |
| Actinobacteriota | 53 | 55 | 108 | 0.50925926 | 16 | 6 |
| Bdellovibrionota_B | 1 | 1 | 2 | 0.5 | 17 | 21 |
| unbinned | 458 | 322 | 780 | 0.41282051 | 18 | 1 |
| Cyanobacteria | 3 | 2 | 5 | 0.4 | 19 | 18 |
| Verrucomicrobiota | 27 | 7 | 34 | 0.20588235 | 20 | 13 |
| Crenarchaeota | 2 | 0 | 2 | 0 | 21 | 21 |
| Patescibacteria | 1 | 0 | 1 | 0 | 21 | 23 |
| Bdellovibrionota | 2 | 0 | 2 | 0 | 21 | 21 |

**Table S9** Rate of novel BGCs by bacterial phylum as determined by min($d_{BiG-FAM}$). Note that UBA8248 is closely related to Tectomicrobia in GTDB and is classified as Tectomicrobia in NCBI.

| ORF | (Source) Definition |
|---|---|
| ctg43_76 | (as5) SMCOG1227:ribosome biogenesis GTP-binding protein YsxC |
| ctg43_79 | (as5) SMCOG1272:TPR repeat-containing protein |
| ctg43_80 | (blastp) hypothetical protein [Acidobacteriota bacterium] |
| ctg43_81 | (blastp) serine O-acetyltransferase [Acidobacteriota bacterium] |
| ctg43_82 | (blastp) DUF1080 domain-containing protein [Acidobacteriota bacterium] |
| ctg43_83 | (blastp) class I SAM-dependent methyltransferase [Acidobacteriota bacterium] |
| ctg43_84 | (as5) SMCOG1013:aminotransferase class-III |
| ctg43_85 | (blastp) hypothetical protein [Acidobacteriota bacterium], closest classified is M20/M25/M40 family metallo-hydrolase [Blastocatellia bacterium] |
| ctg43_86 | (blastp) DNA-3-methyladenine glycosylase [Acidobacteriota bacterium] |
| ctg43_87 | (blastp) DUF1698 domain-containing protein [Acidobacteriota bacterium], methyltransferase domain-containing protein [Acidobacteriota bacterium] |
| ctg43_88 | (blastp) TIM barrel protein [Acidobacteriota bacterium], mannonate dehydratase [Acidobacteriia bacterium] |
| ctg43_89 | (blastp) HAD family phosphatase [Acidobacteriota bacterium] |
| ctg43_90 | (blastp) metallophosphoesterase family protein [Acidobacteriota bacterium] |
| ctg43_91 | (blastp) hypothetical protein [Acidobacteriota bacterium], closest classified is DUF2393 family protein [Acidobacteriia bacterium] |
| ctg43_92 | (blastp) TIGR01458 family HAD-type hydrolase [Acidobacteriota bacterium] |
| ctg43_93 | (as5) NRPS: A-domain plus PP-biniding |
| ctg43_94 | (blastp) isocitrate/isopropylmalate dehydrogenase family protein [Acidobacteriota bacterium] |
| ctg43_95 | (as5) SMCOG1103:FAD dependent oxidoreductase |
| ctg43_96 | (blastp) hypothetical protein [Acidobacteriota bacterium], closest classified is twin-arginine translocation signal domain-containing protein [Acidimicrobiia bacterium] |
| ctg43_97 | (blastp) CehA/McbA family metallohydrolase [Acidobacteriia bacterium] |
| ctg43_98 | (blastp) exo-alpha-sialidase [Acidobacteriota bacterium] |
| ctg43_99 | (blastp) hypothetical protein [Acidobacteriota bacterium], closest classified is family 10 glycosylhydrolase [Acidobacteriota bacterium] |

| | |
|---|---|
| ctg43_103 | (blastp) tetratricopeptide repeat protein [Acidobacteriota bacterium] |
| ctg43_104 | (blastp) HAD family hydrolase [Acidobacteriota bacterium] |
| ctg43_105 | (as5) SMCOG1007:cytochrome P450 |

**Table S10** AntiSMASH (as5) and protein-BLAST (blastp) results for ORFs in c00043 from CS783. Only classifications below an e-value of $e^{-50}$ were recorded.
